# Bacteria detect neutrophils via a system that responds to hypochlorous acid and flow

**DOI:** 10.1101/2022.02.01.478687

**Authors:** Ilona P. Foik, Summer J. Kasallis, Runhang Shu, Serena Abbondante, Michaela E. Marshall, Lauren A. Urban, Leora Duong, Andy P. Huang, Eric Pearlman, Timothy L. Downing, Albert Siryaporn

## Abstract

Neutrophils kill bacteria by producing an array of lethal molecules, including hypochlorous acid (HOCl). The extent to which pathogens detect HOCl from neutrophils and defend themselves has not been understood. We report that the opportunistic pathogen *Pseudomonas aeruginosa* responds to activated neutrophils by upregulating the flow regulated operon (Fro), which was previously shown to be activated by fluid flow. We found that *fro* is upregulated in a mouse infection model where neutrophil influx occurs. This upregulation is induced *in vitro* by HOCl and its secondary product taurine chloramine but not by other neutrophil defense factors including LL-37, histones, or H_2_O_2_. HOCl induces the FroR-dependent upregulation of methionine sulfoxide reductases that relieve otherwise lethal oxidative stress. Fro expression is regulated by FroR’s anti-σ factor FroI, which contains the highest density of methionines and cysteines of all anti-σ factors. The second-order rate constants of HOCl are highest with these residues, raising the possibility that the activation of *fro* could involve oxidation of FroI. These findings suggest a model in which flow transports oxidizing molecules that activate the *fro* operon, establishing an early warning system for *P. aeruginosa* that improves its survival against host immune defenses and persistence during infection.

## INTRODUCTION

The Gram-negative opportunistic pathogen *P. aeruginosa* is a major cause of hospital-acquired infections and is a significant antibiotic resistance threat. To survive in a broad range of conditions, this bacterium has evolved a variety of signaling pathways that detect changes in nutrients, chemical gradients, and flow, and activates responses that promote its survival (Fang et al., 2016; Sanfilippo et al., 2019). Understanding how extracellular-sensing pathways detect and respond to host cues is critical for improving the treatment of bacterial infections and minimizing the generation of antibiotic-resistant bacteria.

Mammalian immune systems respond to the presence of bacterial pathogens by recruiting immune cells that migrate rapidly through the circulatory system to sites of infection (Aymonnier et al., 2023). Neutrophils are the first and most abundant immune cells recruited to inflamed or damaged tissues (Lavoie et al., 2011) and are stimulated into an “activated” state by bacterial components, such as the membrane constituent lipopolysaccharide. Activated neutrophils produce multiple products that are toxic to bacteria including reactive oxygen species (ROS) and reactive chlorine species (RCS) (Hampton et al., 1998; Segal, 2005; Sultana et al., 2020; Ulfig and Leichert, 2020). While the toxic effects of ROS on bacteria are understood, far less is known about how bacteria are affected by RCS (Fang, 2011; Gray et al., 2013). Hypochlorous acid (HOCl) is a major RCS product of neutrophils that is formed through myeloperoxidase enzymes from hydrogen peroxide (H_2_O_2_) and chloride ions, and is strongly bactericidal (Albrich and Hurst, 1982; Harrison and Schultz, 1976; Segal, 2005; Winterbourn et al., 2016). *P. aeruginosa* is a potent activator of neutrophil respiratory bursts (Jensen et al., 1990), which release an abundance of antibacterial components including HOCl, but whether and how the bacteria detect and respond to neutrophil components is not understood.

*P. aeruginosa* responds to shear generated by the flow of fluids via the *fro* operon, which is upregulated in the bacterium during infection in humans (Cornforth et al., 2018; Padron et al., 2023; Sanfilippo et al., 2019) (Figure 1A). In addition, *fro* is necessary for successful colonization of the gastrointestinal tract and lung infections (Potvin et al., 2003; Skurnik et al., 2013), although the reasons for this is unclear. Given its important role in infection, it is possible that Fro could respond to immune-associated cues, such as the presence of neutrophils. The operon consists of *PA14_21570*, *PA14_21580, PA14_21590,* and *PA14_21600*, which was termed *froABCD* (Sanfilippo et al., 2019), and the extracytoplasmic function (ECF) sigma (σ) and anti-σ factors *PA14_21550* and *PA14_21560* (Boechat et al., 2013), which were termed *froR* and *froI* respectively, by (Sanfilippo et al., 2019). Expression of *froABCD* is regulated by FroR and FroI: overexpression of *froR* upregulates *fro,* whereas overexpression of *froI* downregulates *fro* (Sanfilippo et al., 2019). Flow was reported to activate FroR-dependent transcription of *froABCD* through a mechanism that involves chemical transport (Padron et al., 2023).

**Figure 1.**
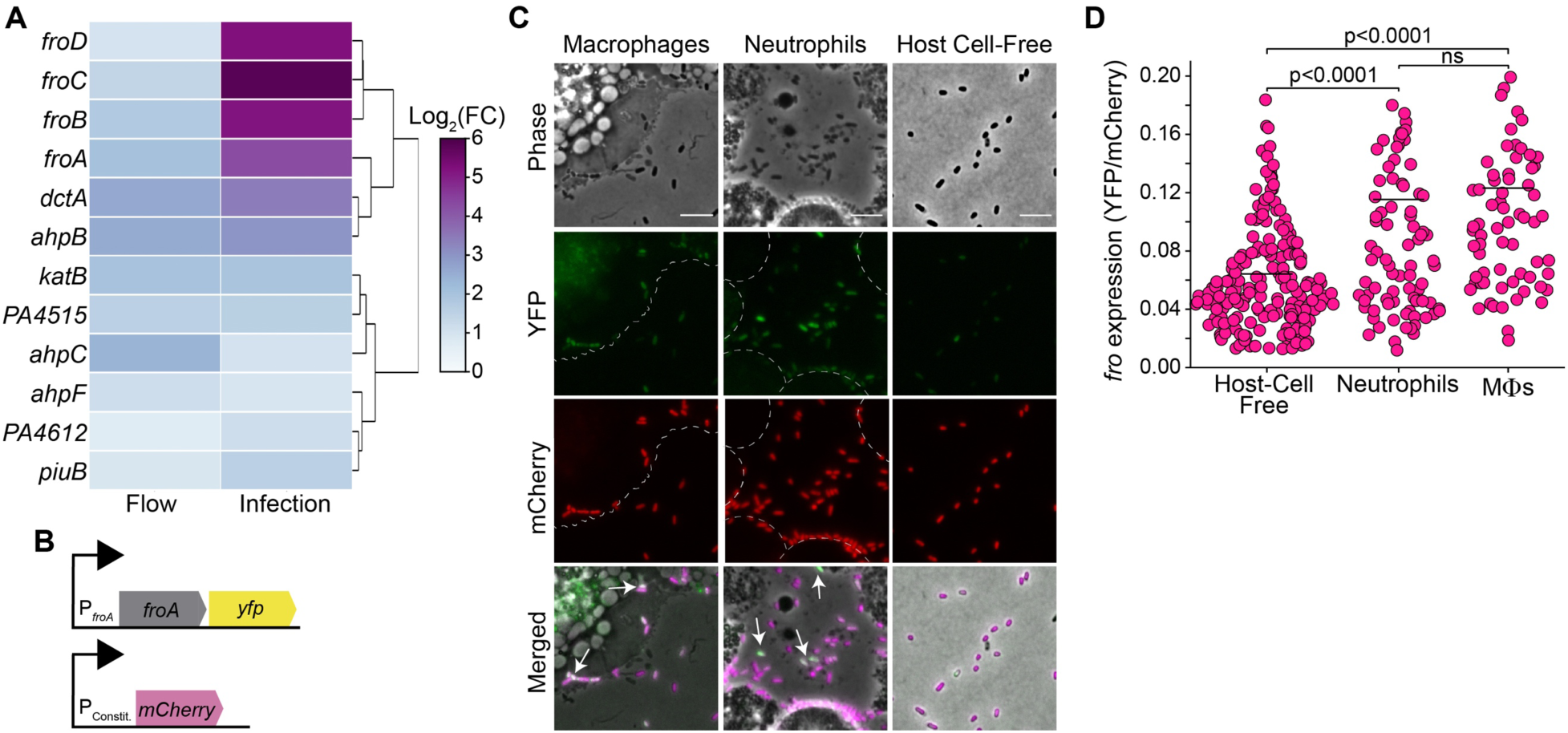
Activation of *fro* promoter transcription by flow and during infection of human tissue. **(A)** RNA-seq heatmap of *P. aeruginosa* genes that are upregulated by at least 3-fold by flow and during infection in human wounds. Datasets are from (Cornforth et al., 2018; Sanfilippo et al., 2019). Log_2_(FC) indicates the log_2_ of the fold-change. **(B)** Schematic of the *P. aeruginosa* fluorescent reporter strain AL143 that contains a transcriptional fusion of *yfp* to the *fro* promoter and expresses mCherry under a constitutive promoter. **(C)** Representative phase-contrast and fluorescence images and **(D)** corresponding *fro* expression as determined by the ratio of YFP to mCherry fluorescence, of *P. aeruginosa* strain AL143 co-incubated with neutrophils, macrophages (MΦs), or host cell-free medium for 30 minutes. Dashed lines in images indicate boundaries of host cells and white arrows indicate *P. aeruginosa* with intense YFP fluorescence relative to that of mCherry. Data points indicate individual *P. aeruginosa* cells from a single experiment and bars indicate the average. Images in (C) are representative of three independent experiments and scale bars represent 5 μm. P-values were obtained using Welch’s t-test of data from single experiments, with values of p>0.05 denoted as ns.

Here, we show that the *fro* operon responds directly to HOCl, which is produced robustly by neutrophils during immune response. Using a reporter system in live bacteria and through quantitative measurements of transcription, we show that *fro* expression is upregulated by activated neutrophils *in vitro* and in a murine model of corneal infection. Through transcriptional profiling, we determine that bacteria mitigate the toxic effects of HOCl through a FroR-dependent mechanism that activates methionine sulfoxide reductases to relieve oxidative stress. These data suggest a novel mechanism in which bacterial defense against host innate immunity is primed by a single system that detects both flow and HOCl.

## RESULTS

### Macrophages and neutrophils activate *fro* expression in *P. aeruginosa* under non-flow conditions

To investigate whether immune cells directly activate the *fro* operon, we quantified *fro* expression using a *P. aeruginosa* strain that co-expresses the yellow fluorescent protein (YFP) under the transcriptional control of the *fro* promoter and mCherry under the control of a constitutive promoter (Figure 1B). While *fro* expression increased under flow (Figure 1—figure supplement 1A), experiments in this study were performed in the absence of flow in order to deconvolve the effects of flow from potential activation of *fro* by immune cells. Because neutrophils and macrophages are the first immune cells recruited to sites of infection, we co-incubated *P. aeruginosa* with human neutrophils or mouse macrophages for 30 minutes. Relative to *P. aeruginosa* that were incubated in host cell-free media, *P. aeruginosa* co-incubated with either neutrophils or macrophages increased YFP/mCherry ratios by comparable levels (Figure 1C-D), suggesting that close proximity to or direct interaction with these host immune cells increases *fro* expression in *P. aeruginosa*.

### Activated neutrophils induce *fro* expression

To assess whether *fro* upregulation in *P. aeruginosa* could be part of a response to respiratory bursts from neutrophils and macrophages, we focused on the *fro* response to neutrophils (Figure 2A) because these cells produce larger respiratory bursts and greater levels of ROS than macrophages (Colombo et al., 2017; Forman and Torres, 2002; Thekkan et al., 2019). *P. aeruginosa* were incubated in conditioned medium from human neutrophils that were stimulated using phorbol 12-myristate 13-acetate (PMA), which triggers respiratory bursts (Karlsson et al., 2000). The expression of *fro* was quantified by averaging the ratios of YFP to mCherry fluorescence for at least one hundred individual *P. aeruginosa* cells. In support of our hypothesis, conditioned medium from PMA-stimulated neutrophils significantly increased *fro* expression in bacteria compared to conditioned medium from unstimulated neutrophils (Figure 2B-D, Figure 2—source data 1 and 2). We confirmed that this *fro* response was not due to PMA alone, as PMA-treated medium without neutrophils had no effect on *fro* expression (Figure 2B-C). Similarly, conditioned medium from unstimulated neutrophils had no significant effect on *fro* expression compared to medium alone, suggesting that the upregulation of *fro* is due to respiratory burst products.

**Figure 2.**
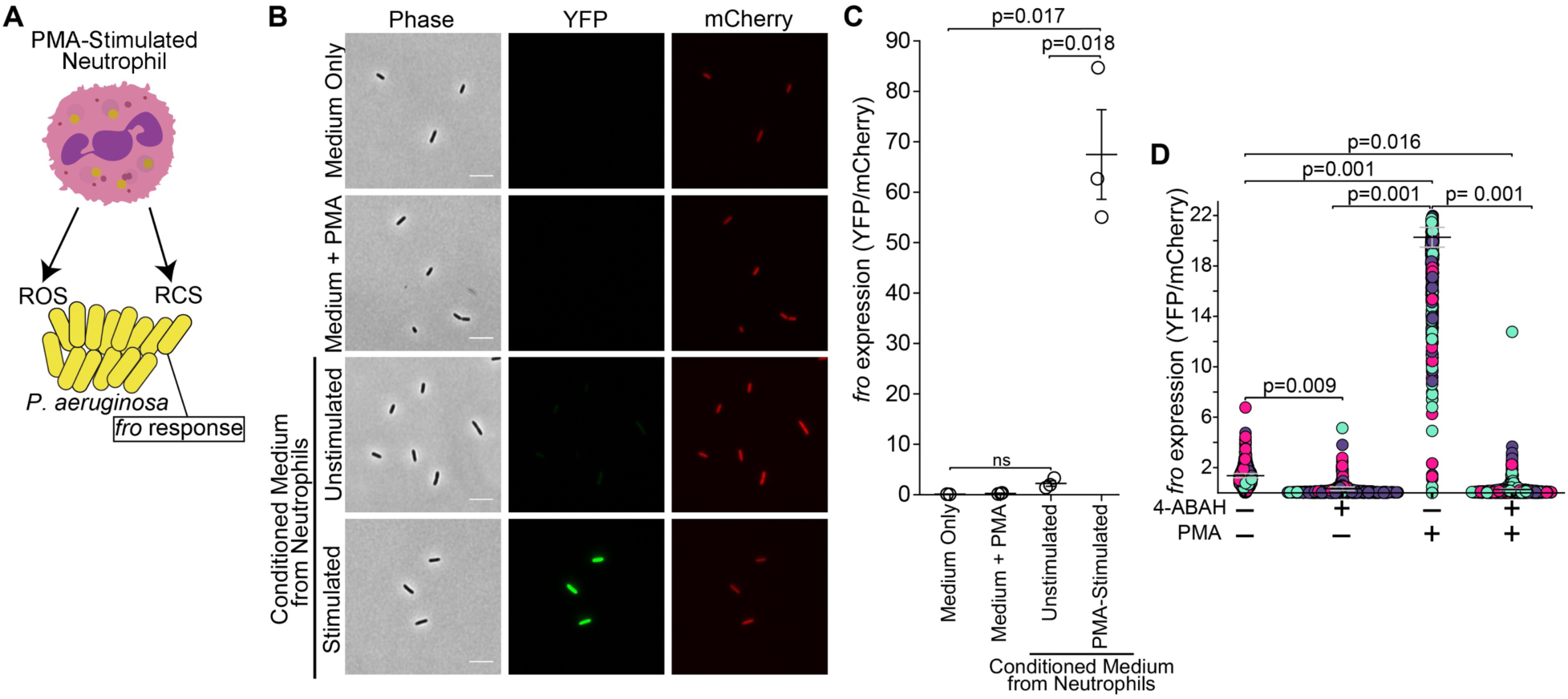
Conditioned medium from stimulated neutrophils induces *fro* expression. **(A)** Schematic depicting that PMA-stimulated neutrophils produce respiratory burst products that could activate *fro* expression. **(B)** Representative phase-contrast and fluorescence images and **(C)** *fro* expression as determined by ratios of YFP to mCherry fluorescence, of *P. aeruginosa* strain AL143 cultured for 3 hours in medium only, medium with PMA, conditioned medium from PMA-stimulated neutrophils, or conditioned medium from unstimulated neutrophils. Scale bars represent 5 μm. Data points indicate the average from at least one hundred individual *P. aeruginosa* cells from a single experiment and horizontal bars indicate the mean of at least three independent experiments. **(D)** *fro* expression as determined by ratios of YFP to mCherry fluorescence of *P. aeruginosa* strain AL143 that was cultured for 3 hours in conditioned medium in which neutrophils were stimulated with 100 ng/mL PMA, pre-treated with 2mM 4-ABAH, or both. Data points indicate individual *P. aeruginosa* cells, with each color representing an independent experiment, bars represent the average of 3 independent experiments, each using separate neutrophil isolations, and error bars indicate standard error of the mean (SEM). P-values were obtained using Welch’s t-test, with values of p>0.05 denoted as ns.

HOCl and other RCS are produced from ROS during respiratory bursts via myeloperoxidase (MPO). To investigate whether *P. aeruginosa fro* expression is induced by MPO-derived RCS, we pre-treated neutrophils prior to PMA stimulation with 4-aminobenzoic acid hydrazide (4-ABAH) (Figure 2D), a specific and irreversible MPO inhibitor (Kettle et al., 1997). Indeed, the activation of *fro* expression by PMA-stimulated neutrophil medium was abrogated in neutrophils that were pre-treated with the MPO inhibitor. The suppression of *fro* activation by the 4-ABAH pre-treatment of neutrophils prior to PMA-stimulation followed a dose-dependent trend (Figure 2—figure supplement 1). This result is consistent with a graded *fro* response to decreasing RCS concentrations as a result of increasing 4-ABAH. HOCl production in neutrophils due to PMA stimulation and its inhibition by 4-ABAH was confirmed using a fluorometric HOCl-specific assay (Figure 2—figure supplement 2). These results indicate that *fro* upregulation depends on MPO activity and suggest that *fro* may be activated in response to HOCl.

### HOCl but neither H_2_O_2_ nor HNO_3_ induces *fro* expression

ROS and RCS have inhibitory effects on bacterial growth. To explore the hypothesis that these classes of molecules induce *fro* expression in *P. aeruginosa*, we measured the effects of H_2_O_2_, HNO_3_, and NaOCl on *fro* expression at concentrations approaching their minimum inhibitory concentrations (MICs) (Figure 3A). NaOCl sharply increased *fro* expression between 0.25 to 1 µM (Figure 3B, Figure 3—figure supplement 1). To validate that the increase in YFP intensity was due to increased transcription, reverse transcription quantitative PCR (RT-qPCR) was also performed, which yielded a 200-fold increase in *fro* transcription in response to 1 µM NaOCl treatment (Figure 3C). While 2 µM NaOCl caused a relatively small increase in YFP expression (Figure 3B and Figure 3—figure supplement 1), we attribute the diminished expression to the quenching of YFP by oxidative species, to which YFP is particularly sensitive (Tsourkas et al., 2005), and to the effects of NaOCl on cell viability, as this concentration also decreased cell size (Figure 3—figure supplement 1).

**Figure 3.**
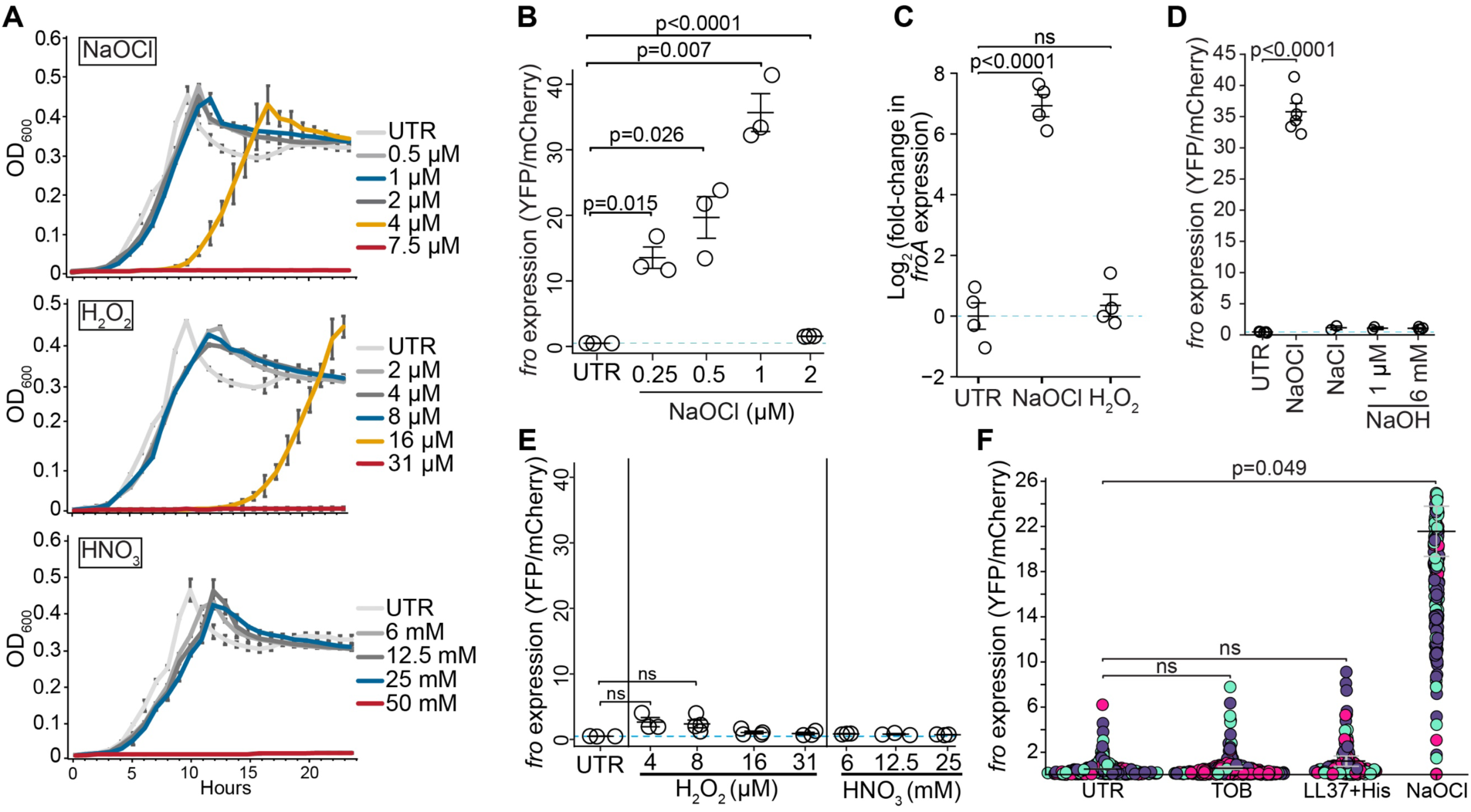
NaOCl but neither H_2_O_2_ nor HNO_3_ induces *fro* expression. **(A)** Growth profiles as measured by optical density (OD_600_) of *P. aeruginosa* treated with the indicated concentrations of NaOCl, H_2_O_2_, HNO_3_, or no treatment (UTR). Data points indicate a mean of at least two independent experiments and error bars indicate SEM. **(B)** *fro* expression as determined by ratios of YFP to mCherry fluorescence in *P. aeruginosa* after 3 hour treatment with the indicated concentrations of NaOCl or no treatment (UTR). **(C)** Abundance of *fro*A transcripts in *P. aeruginosa* after 30 minutes of treatment with 1 µM NaOCl, 4 µM H_2_O_2_, or no treatment (UTR), as measured by RT-qPCR. Measurements were normalized to 5S ribosomal RNA and the logarithm (base 2) of the fold-change in transcription was computed relative to the untreated condition. **(D and E)** *fro* expression as determined by ratios of YFP to mCherry fluorescence in *P. aeruginosa* after 3 hour treatment with **(D)** 1 µM NaOCl, 1 µM NaCl, 1 µM or 6 mM NaOH, or no treatment, and **(E)** indicated concentrations of H_2_O_2_, HNO_3_, or no treatment (data replotted from panel B). In panels B, D, and E, data points indicate the average from at least one hundred individual *P. aeruginosa* cells in each experiment. **(F)** *fro* expression as determined by ratios of YFP to mCherry fluorescence of *P. aeruginosa* after 3 hour treatment with vehicle control (UTR; sterile water), 0.5 µg/mL tobramycin (TOB), 40 µg/mL LL-37 with 10 µg/mL histones (LL37+His), or 1 µM NaOCl. Data points indicate individual *P. aeruginosa* cells, and different colored data points represent independent experiments. In panels B-F, horizontal bars indicate the mean of independent experiments, error bars indicate SEM, and blue dashed lines indicate the average of the untreated condition. All data were acquired using the *P. aeruginosa* strain AL143, and p-values were obtained using Welch’s t-test with values of p>0.05 denoted as ns.

To rule out that the individual sodium or chloride ion components of NaOCl were responsible for the increase in *fro* expression, we repeated experiments using NaCl, which yielded no change in *fro* expression (Figure 3D). To assess whether the *fro* response may be affected by pH, we tested NaOH at the same concentration as NaOCl, and at a concentration at which a change in pH could be discerned. Due to the robust buffering capacity of the medium, a pH change of ∼0.1 required 6 mM NaOH, which is a concentration that had no obvious effect on growth but is significantly higher than the concentration of NaOCl needed to activate *fro* expression. Regardless, neither concentration of NaOH had any significant impact on *fro* expression (Figure 3D). As *fro* activation is not due to sodium or chloride ions nor changes in pH, we attributed the effects to hypochlorite (OCl^-^), which is in equilibrium with hypochlorous acid (HOCl) at the pH of 7.1 in the medium used in our experiments.

To address the possibility that *fro* expression is activated by oxidizers broadly, we quantified *fro* expression in response to H_2_O_2_ and HNO_3_, at concentrations up to their respective MICs (Figure 3A). While both are strong oxidizers, HNO_3_ is not produced by neutrophils. Neither H_2_O_2_ nor HNO_3_ induced any significant changes in *fro* expression up to near-MIC concentrations (Figure 3E). The lack of *fro* expression due to H_2_O_2_ treatment was not due to defects in fluorescent protein production, as H_2_O_2_ did not induce any change in *fro* transcript abundance, as assayed using RT-qPCR (Figure 3C). Together, these indicate that strong oxidation alone is not sufficient to upregulate *fro*.

We assessed whether other antimicrobial components of neutrophils or other bacterial stressors could be responsible for the *fro* response. Stimulated neutrophils release the antimicrobial peptide LL-37 and histones, which together constitute a synergistic antimicrobial attack (Doolin et al., 2020; Duong et al., 2025). Tobramycin is an aminoglycoside, a class of antibiotics that induces oxidative stress within bacteria (Kohanski et al., 2008). Neither the combination of LL-37 and histones nor tobramycin at near-MIC concentrations induced *fro* expression (Figure 3F). Together, these results indicate that *fro* expression is activated specifically by the neutrophil effector HOCl and not by intracellular oxidative stress or other neutrophil defense factors including LL-37, histones, or H_2_O_2_.

### Corneal infection activates *fro* expression in bacteria

*P. aeruginosa* is a major cause of corneal ulcers and keratitis, which can cause visual impairment and blindness. Neutrophils are recruited within hours to corneal epithelial abrasions, with neutrophil cell density peaking at 18 hours (Li et al., 2006). We tested the hypothesis that *fro* expression is upregulated during corneal infection using a murine bacterial keratitis model in which *P. aeruginosa* strain PAO1F is inoculated onto a corneal abrasion, which we previously demonstrated results in significant neutrophil recruitment to the cornea (Minns et al., 2023; Ratitong et al., 2022). Mice corneas were harvested 0, 2, or 20 hours post-infection and *fro* transcripts were quantified using RT-qPCR. Transcript abundance was normalized to the number of *P. aeruginosa* cells using 5S ribosomal RNA. In support of the hypothesis, *fro* expression increased significantly by 20 hours post-infection (Figure 4). Expression levels showed no significant change within the first two hours of infection, which we attribute to a lower level of immune cell recruitment during this period. The increase in *fro* expression 20 hours post-infection was not due to differences between the *P. aeruginosa* strains, because *fro* transcription in PAO1F also increased in response to NaOCl but not H_2_O_2_ (Figure 4—figure supplement 1), consistent with previous results.

**Figure 4.**
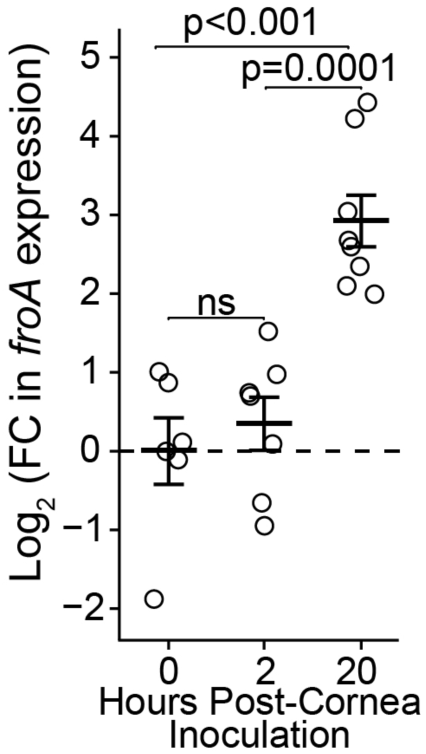
The expression of fro is activated during corneal infection. Abundance of *fro*A transcripts in *P. aeruginosa* strain PAO1F as measured by RT-qPCR, at 0, 2, or 20 hours post-inoculation onto abraded mouse corneas. Measurements were normalized to 5S ribosomal RNA and the logarithm (base 2) of the fold-change in transcription was computed relative to the initial inoculum (time=0 hrs). Log_2_(FC) indicates the log_2_ of the fold-change. Data points represent independent corneal infections, horizontal bars indicate the average of at least six independent experiments, error bars represent SEM, and dashed line indicates the average of the 0h condition. P-values were obtained using Welch’s t-test, with values of p>0.05 denoted as ns.

### Methionine sulfoxide reductase upregulation requires FroR

FroR is the σ factor required for activation of the *fro* operon in response to flow (Sanfilippo et al., 2019). To understand the response to HOCl, we performed transcriptional profiling of wild-type (WT) and Δ*froR P. aeruginosa* strains. The genes that were most upregulated by NaOCl in the WT strain compared to the Δ*froR* mutant were those in the *fro* operon, the phosphate and pyrophosphate-specific outer membrane porin genes *oprO* and *oprP,* and the methionine sulfoxide reductase genes *msrB, msrP,* and *msrQ* (Figure 5A and Appendix 1—table 1). Methionine sulfoxide reductases (MSRs) have a critical role in bacterial survival and defense against RCS stress. HOCl oxidizes methionine and cysteine residues 100-fold more rapidly than other amino acids (Pattison and Davies, 2006; Winterbourn, 1985) and oxidation of methionine in bacterial membrane proteins by neutrophil-produced HOCl correlates with increased bactericidal killing of *E. coli* and *S. aureus* (Rosen et al., 2009). HOCl oxidation of methionine forms methionine sulfoxide, which impairs protein function. MSRs restore proper protein function by reducing methionine sulfoxide back to methionine (Chandra et al., 2023; Gray et al., 2013). Our data thus suggest that FroR is required for the upregulation of MSRs in response to HOCl stress. We did not observe upregulation of PA14_27570, which is homologous to MsrC and reduces strictly free methionine sulfoxide, or of PA14_66330 (MsrA), which reduces both free and protein-bound methionine sulfoxide (Appendix 1—table 1). No significant changes in the expression of enzymes that repair cysteine oxidation, such as thioredoxin, glutathione redoxin, or disulfide repair, were observed, suggesting that the *fro* response is directed towards repairing methionine oxidation.

**Figure 5.**
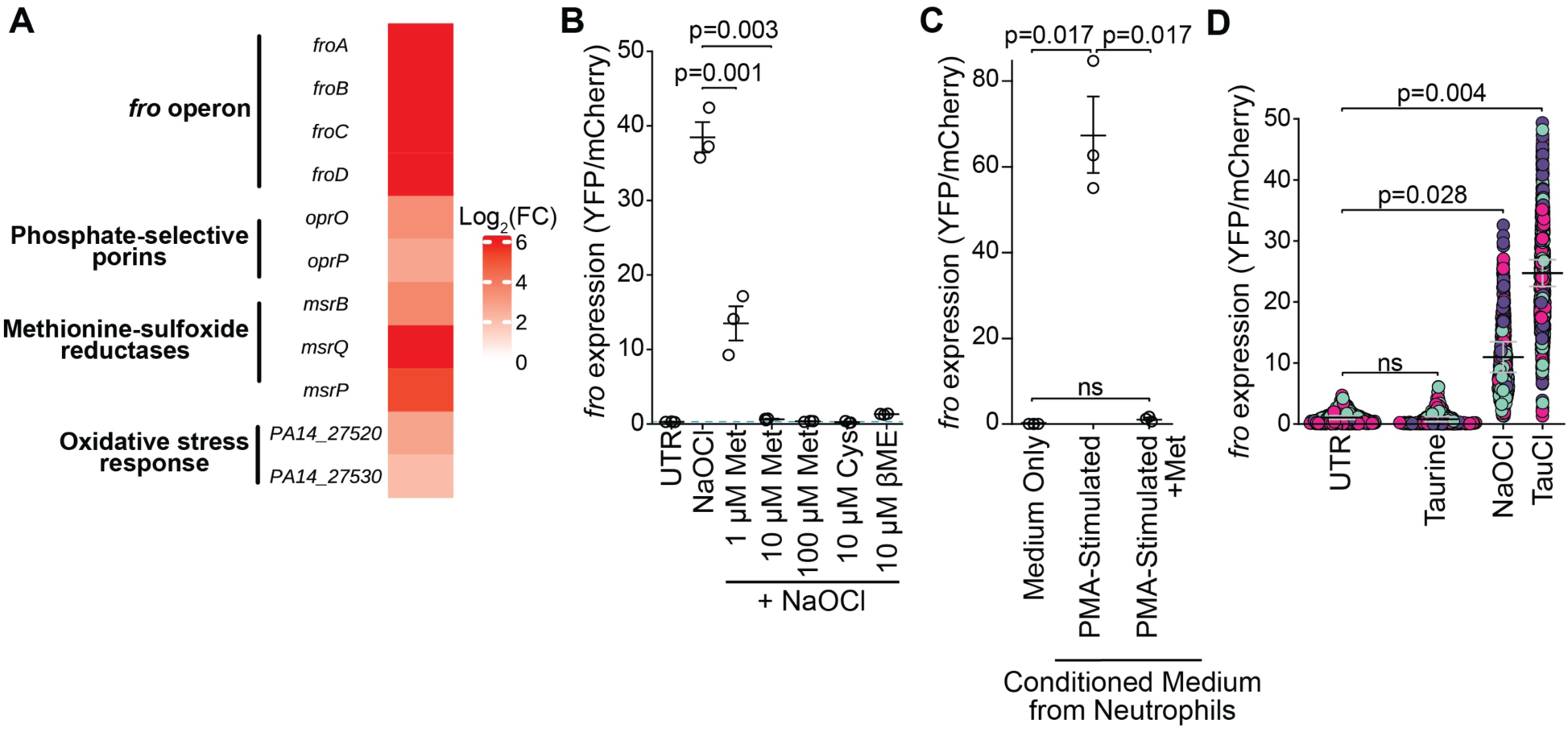
Expression of *fro* is inhibited by methionine and antioxidants and is activated by the secondary HOCl product taurine chloramine. **(A)** Heatmap showing *P. aeruginosa* gene transcripts (sorted by genomic locus) that were upregulated by at least 4-fold in the wild type PA14 background (strain AL143) compared to the Δ*froR* mutant (AL325) after 30 minutes of treatment with 1 µM NaOCl and that had p-values less than 0.05 (n≥3). Log_2_(FC) indicates the log_2_ of the fold-change. Raw data can be found online (Foik, Ilona P. et al., 2025) in NCBI’s Gene Expression Omnibus (see Key Resources Table). (B, C, and D) *fro* expression as determined by ratios of YFP to mCherry fluorescence in strain AL143 after 3 hour **(B)** treatment with vehicle control (UTR), 1 µM NaOCl, or 1 µM NaOCl with combined with methionine (Met), cysteine (Cys), or β-mercaptoethanol (βME) at the indicated concentrations; **(C)** incubation in medium only (replotted from Figure 2C), or conditioned medium from stimulated neutrophils without or with 100 µM methionine (+Met); or **(D)** treatment with vehicle control (UTR; reaction buffer), 1 µM taurine, 1 µM NaOCl, or 1 µM taurine chloramine (TauCl). Data points in **B** and **C** indicate the average from at least one hundred individual *P. aeruginosa* cells or in **D**, individual *P. aeruginosa* cells with different colors representing independent experiments. Blue dashed line in panel B indicates the average of the untreated condition. Horizontal bars indicate the mean of at least three independent experiments and error bars represent SEM. P-values were obtained using Welch’s t-test, with values of p>0.05 denoted as ns.

We reasoned that HOCl could oxidize methionine residues within a *P. aeruginosa* sensor, which could in-turn upregulate *fro* expression. A number of anti-σ factors contain sensory domains that transduce sensory signals to their cytoplasmic σ factor (Paget, 2015). FroI is an anti-σ factor for the FroR σ factor (Boechat et al., 2013), which raises the possibility that HOCl or its secondary products could oxidize FroI directly, resulting in the activation of FroR. Consistent with this possibility, we found that FroI contains the highest concentration of methionine and cysteine residues of all identified *P. aeruginosa* PA14 anti-σ factors (Appendix 2—table 1).

We hypothesized that *fro* activation would be suppressed in the presence of excess of free methionine, as it would reduce available HOCl and thus oxidation of the FroI sensor. Indeed, co-treatment of NaOCl with methionine suppressed *fro* expression (Figure 5B). We reasoned that other antioxidants such as cysteine and β-mercaptoethanol (βME) could also function as alternative oxidation targets for HOCl, and therefore also suppress *fro*. Consistent with this hypothesis, co-treatment of NaOCl with cysteine or βME significantly suppressed *fro* expression compared to cysteine or βME alone (Figure 5B and Figure 5—figure supplement 1A-B).

We considered how *fro* could be upregulated by stimulated neutrophils in the context of these findings. HOCl and other reactive chlorine species produced by neutrophils could oxidize the *P. aeruginosa* sensor, resulting in *fro* upregulation. Under this interpretation, supplying excess methionine to conditioned medium from activated neutrophils, which would reduce HOCl, should also inhibit *fro* upregulation. Indeed, supplementing with methionine completely suppressed *fro* expression (Figure 5C). Together, these data suggest a model in which *fro* responds to HOCl-induced oxidation of a sensor by upregulating MSRs.

While our data shows that HOCl can directly activate *fro* expression, HOCl reacts with neutrophil components, forming secondary RCS, including monochloramines, protein-derived chloramines, and taurine chloramine (Kim and Cha, 2014; Klebanoff et al., 2013; Ulfig and Leichert, 2020). Taurine chloramine is an expected major RCS released from neutrophils due to the high abundance of taurine in these cells (Kim and Cha, 2014; Marcinkiewicz and Kontny, 2014) and retains oxidizing potential towards cysteine and methionine (Peskin and Winterbourn, 2001). Indeed, taurine chloramine directly activated *fro* expression (Figure 5D).

### FroR protects *P. aeruginosa* against HOCl

The upregulation of MSRs is expected to improve bacterial growth against HOCl stress. We measured the growth profiles of WT and Δ*froR* strains of *P. aeruginosa*. Treatment with 4 µM NaOCl significantly inhibited the growth of the WT strain (Figure 6A) but resulted in even greater growth inhibition in the Δ*froR* strain. This observation suggests that the *fro* operon improves *P. aeruginosa* tolerance to HOCl stress. We note that diminished *fro* expression was observed at 2 µM NaOCl compared to 1 µM in the YFP reporter strain (Figure 3B). It is possible that this reduction is due to YFP oxidation and that *fro* operon components are expressed at higher concentrations of NaOCl.

**Figure 6.**
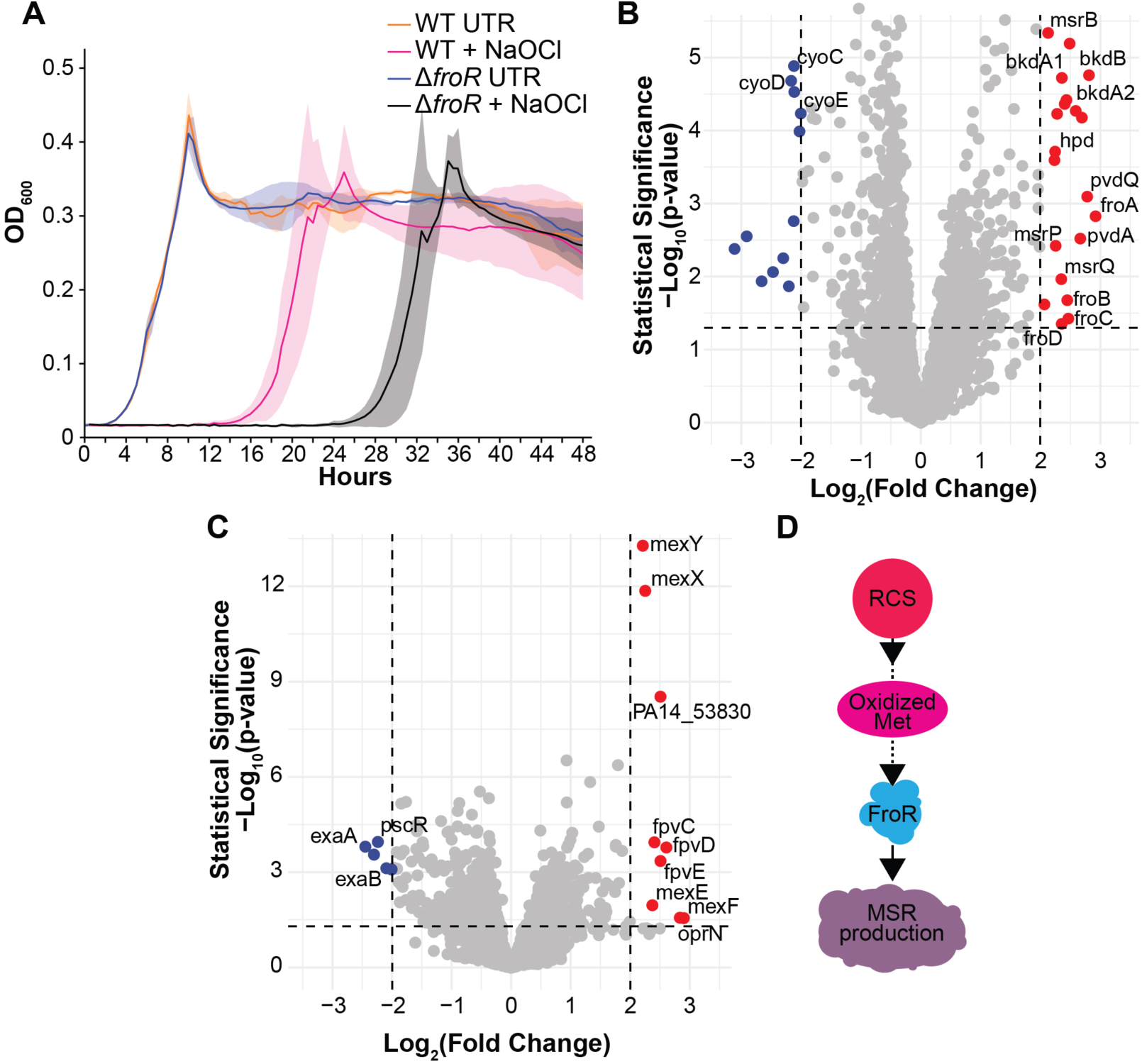
FroR improves P. aeruginosa growth against HOCl stress and regulates transcription of genes involved in antioxidant defense. **(A)** Growth profiles of wild-type (WT) *P. aeruginosa* strain AL143 and the Δ*froR* mutant strain AL325 treated with 4 µM NaOCl or vehicle control (UTR; sterile water). Data points represent an average of at least three independent experiments and error bars indicate SEM. **(B-C)** Statistical significance as a function of fold-change in transcript abundance in **(B)** WT and **(C)** Δ*froR* after treatment with 1 µM NaOCl (n≥3), as determined by RNA sequencing. Genes in red (upregulated) and blue (downregulated) were altered by at least four-fold. The unlabeled gene loci shown have not been previously annotated. P-values were determined using the Wald test (Love et al., 2014). Greater values along the vertical axis indicate greater statistical significance and the dashed horizontal line indicates a p-value of 0.05. Raw data can be found online (Foik, Ilona P. et al., 2025) in NCBI’s Gene Expression Omnibus (see Key Resources Table). **(D)** Schematic that depicts a model in which RCS oxidizes methionine residues (Met) in a target sensor(s), such as the ECF anti-σ factor FroI, which in turn activates the FroR σ factor and increases the production of methionine sulfoxide reductases (MSR).

We considered other genes which may improve WT *P. aeruginosa* tolerance to HOCl. HOCl increased the expression of iron uptake genes (*pvdA* and *pvdQ*) and downregulated the expression of oxidative phosphorylation genes (*cyoCDE*) (Figure 6B and Appendix 1—table 2). These changes suggest that HOCl causes an iron deficiency and decreases oxidative phosphorylation. In order to compensate for decreased oxidative phosphorylation, *P. aeruginosa* may utilize amino acid catabolism as an alternative energy source, as evidenced by a concomitant upregulation of amino acid catabolism genes (*hpd, bkdA1, bkdA2,* and *bkdB*) by HOCl (Figure 6B and Appendix 1—table 2).

While the Δ*froR* mutant was more sensitive to HOCl than WT, its growth was not fully inhibited (Figure 6A), suggesting that additional mechanisms relieve HOCl stress. In the Δ*froR* mutant, HOCl upregulated genes encoding multidrug efflux pumps (*mexEF-oprN* and *mexXY*) (Figure 6C and Appendix 1—table 3). These pumps can relieve oxidative stress (Fargier et al., 2012; Fraud and Poole, 2011) but were not activated in the WT strain. This observation suggests that bacterial defense against HOCl may be hierarchical, in which Fro-regulated and Mex pathways could be first-line and second-line responses to HOCl stress, respectively. Supporting this interpretation, it has been reported that treatment of the WT strain using significantly higher (millimolar) HOCl concentrations upregulates *mexEF*-*oprN* (Farrant et al., 2020) and the *mexXY* regulator *mexT* (Groitl et al., 2017).

## DISCUSSION

Neutrophils are activated by the presence of bacteria that infect host tissue. In their activated state, neutrophils generate respiratory bursts that produce an abundance of HOCl. We have demonstrated that *P. aeruginosa* responds to activated neutrophils by upregulating the transcriptional expression of the *fro* operon, which relieves HOCl stress. Our data suggest that the upregulation of *fro* could function as a bacterial defense mechanism against neutrophil attack, thus improving *P. aeruginosa* survival and colonization of human tissue during infection.

### Detection and response to HOCl

Based on (i) our observation that the expression of multi-drug efflux pumps is activated in the absence of FroR (Figure 6C and Appendix 1—table 3), (ii) that this expression occurs at high HOCl concentrations (Farrant et al., 2020), and (iii) that multiple systems repair HOCl damage, the HOCl response could follow a hierarchy of activation. FroR activates two distinct classes of reductases that repair HOCl-oxidized methionine in proteins: MsrB reduces methionine sulfoxide in the cytoplasm using electrons provided by the thioredoxin system (Ezraty et al., 2005; Lourenço Dos Santos et al., 2018), while MsrPQ works in the periplasm and inner membrane using electrons from the respiratory chain (Gennaris et al., 2015). The activation of these two reductase types enables *P. aeruginosa* to limit HOCl stress in separate cellular spaces by utilizing distinct electron donor sources.

How could *P. aeruginosa* detect HOCl? We speculate that as an anti-σ factor, FroI could immobilize FroR, inhibiting FroR from reaching transcriptional targets. Given the particularly high density of methionine and cysteine residues in the FroI, we propose that this protein could serve as an HOCl sensor by releasing the sigma factor FroR in response to oxidation of these residues (Figure 6D). Given that taurine chloramine activates *fro* expression, it is possible that taurine chloramine or other secondary RCS could oxidize FroI. Future work will need to assess the impact of oxidation on FroI and its association with FroR.

### Relationship with neutrophil-mediated killing and flow

One of the striking features of the Fro system is that it is tuned for maximal activation at micromolar HOCl concentrations that are close to the MIC. While our experiments measured the effects of HOCl at steady-state using medium that does not contain amino acids, host proteins, and other host factors, HOCl dynamics in host environments are much more complex, where HOCl production is estimated to be in quantities that exceed the sensitivity of Fro. For example, HOCl production is predicted in excess of 100 mM per minute in the phagosome (Winterbourn et al., 2006). In addition, stimulated neutrophils at densities comparable to those found in blood (Tahir and Zahra, 2026) using PMA produces extracellular HOCl at 80 µM per hour, as measured through taurine chloramine absorbance (Weiss et al., 1982). Additional measurements using slight changes in stimulation time, neutrophil density, or PMA concentration yielded comparable HOCl production rates within a factor of 2 (Dypbukt et al., 2005; Test et al., 1984). At sites of inflammation, HOCl production in interstitial fluid has been predicted to be in the millimolar range due to the greater density of neutrophils there (Weiss, 1989).

Despite the high production rates of HOCl, the steady-state concentration in neutrophils and *in vivo* is expected to be significantly lower due to its rapid conversion into secondary RCS through reaction with neutrophil or plasma components. For example, the lifetime of HOCl is estimated to be 0.1 µs and have an action radius under 0.1 µm in the phagosome (Schürmann et al., 2017), which would mitigate the antimicrobial effects of HOCl but retain oxidizing potential. The secondary RCS taurine chloramine is long-lived, well-tolerated in the body, and retains significant bactericidal activity in the micromolar range (Gottardi and Nagl, 2010; Kim and Cha, 2014; Nagl et al., 2000). It is plausible that diminished HOCl or longer-lived secondary RCS fall within the activating range of Fro, which would in turn mitigate their antimicrobial effects through upregulation of the MSRs. Importantly, our observation that *fro* expression is activated during corneal infection, which involves significant neutrophil recruitment, supports a role for the Fro response in a neutrophil-dense environment. In addition, *fro* expression was activated by macrophages, which can also contain MPO—albeit much less (Odobasic et al., 2016), suggesting that macrophage MPO-produced HOCl or another macrophage-produced species could also activate *fro*.

An additional layer of complexity of understanding the role of Fro during infection is that flow exposes bacteria to higher concentrations of molecules in a fluid. The bacterial response to flow, which was initially referred to as rheosensing (Sanfilippo et al., 2019), was more recently attributed to the effects of molecular transport by flow (Padron et al., 2023). In particular, flow stimulated a response to H_2_O_2_ in cells that were otherwise unresponsive to it in the absence of flow (Padron et al., 2023). Flow could similarly amplify the stimulating effects of RCS, causing *P. aeruginosa* to respond to circulating neutrophils that produce RCS at much lower concentrations. At the same time, flow could amplify the bactericidal effects against *P. aeruginosa*. Future work will need to determine the relative steady-state abundances of different secondary RCS and address their impact in combination with flow on bactericidal activity and the activation of the Fro response.

At the fundamental level, Fro is a system that increases antimicrobial tolerance, which can promote antimicrobial resistance. For example, bacteria that are exposed to low concentrations of HOCl have higher rates of horizontal gene transfer and can acquire antibiotic resistance genes more rapidly (Hou et al., 2019; Jin et al., 2020). The Fro system could thus facilitate antibiotic resistance through increasing *P. aeruginosa* tolerance to HOCl. Our findings suggest that the bacterial HOCl response could thus be an important bacterial defense mechanism to target in the development of novel antibiotic therapeutics. Inhibition of this pathway could increase *P. aeruginosa* susceptibility to neutrophil-mediated killing while minimizing antibiotic resistance.

## MATERIALS AND METHODS

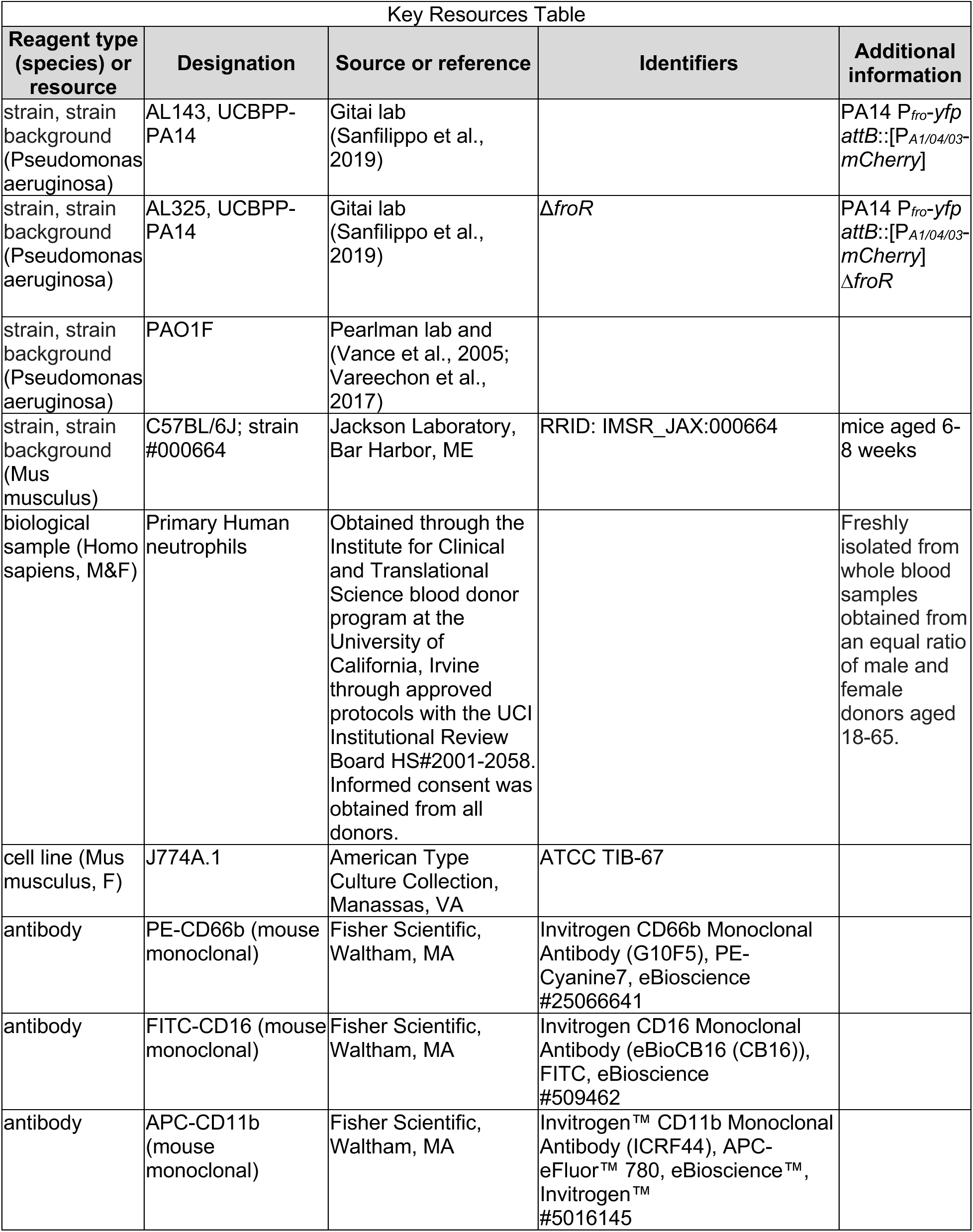
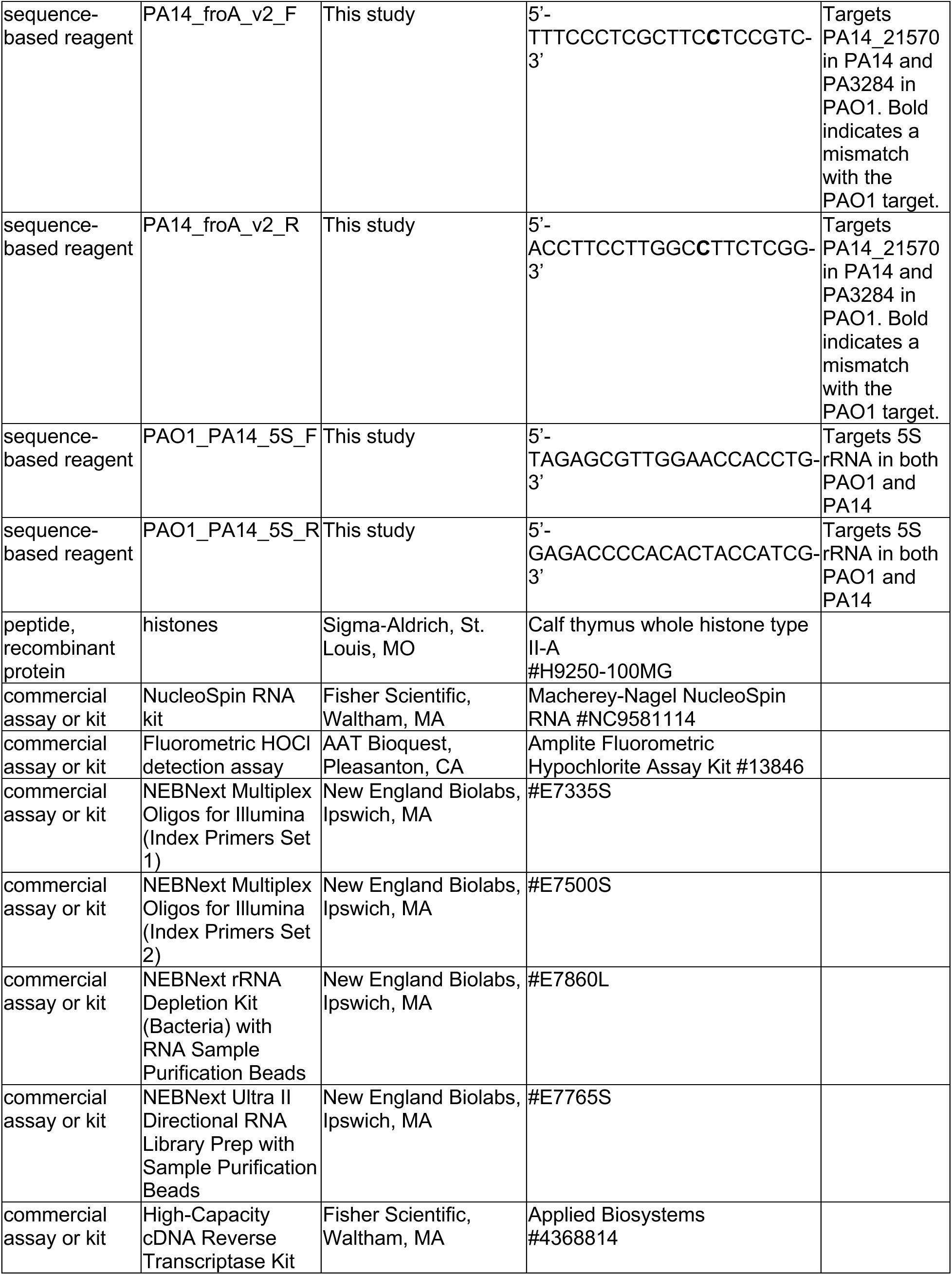
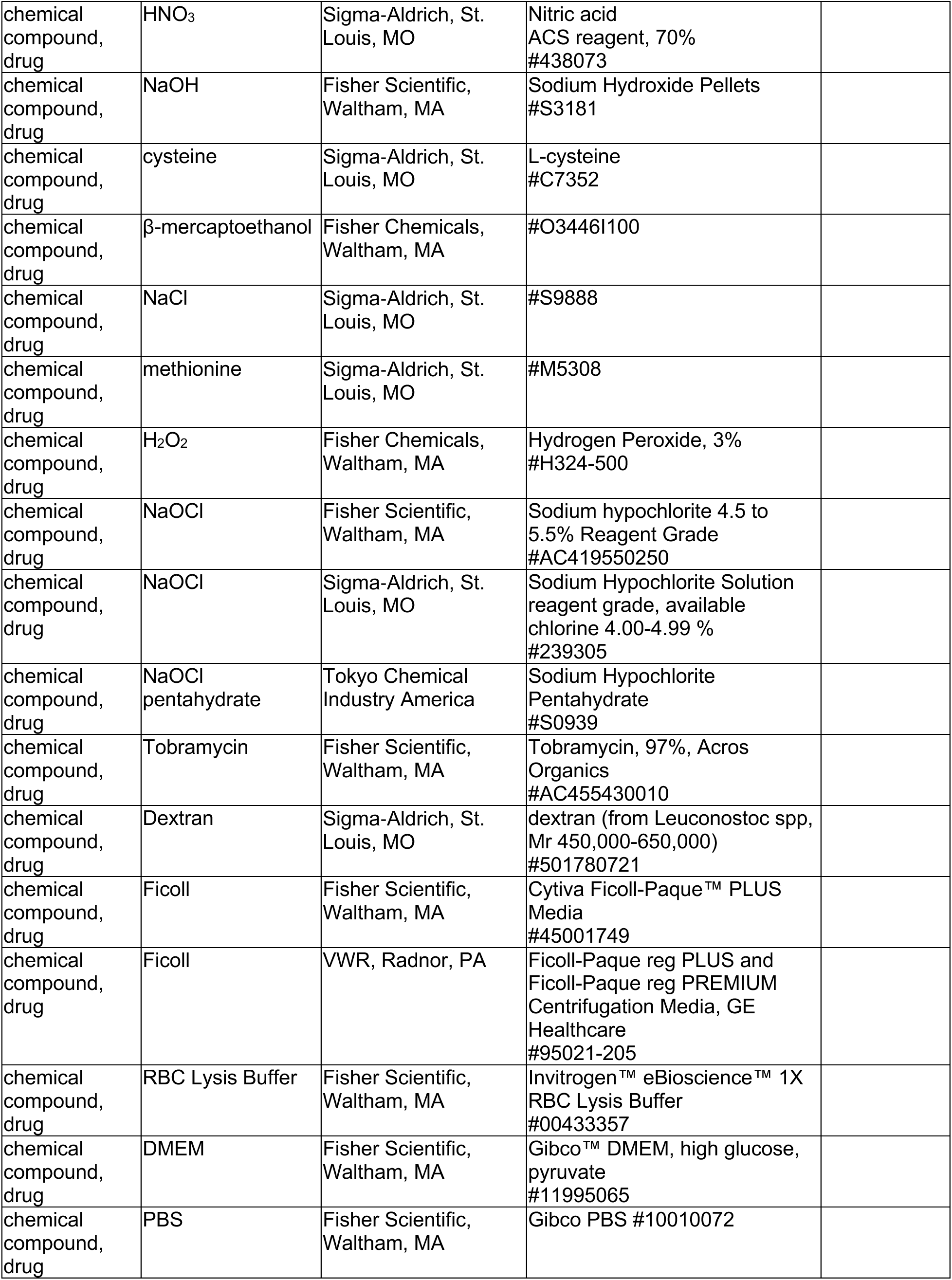
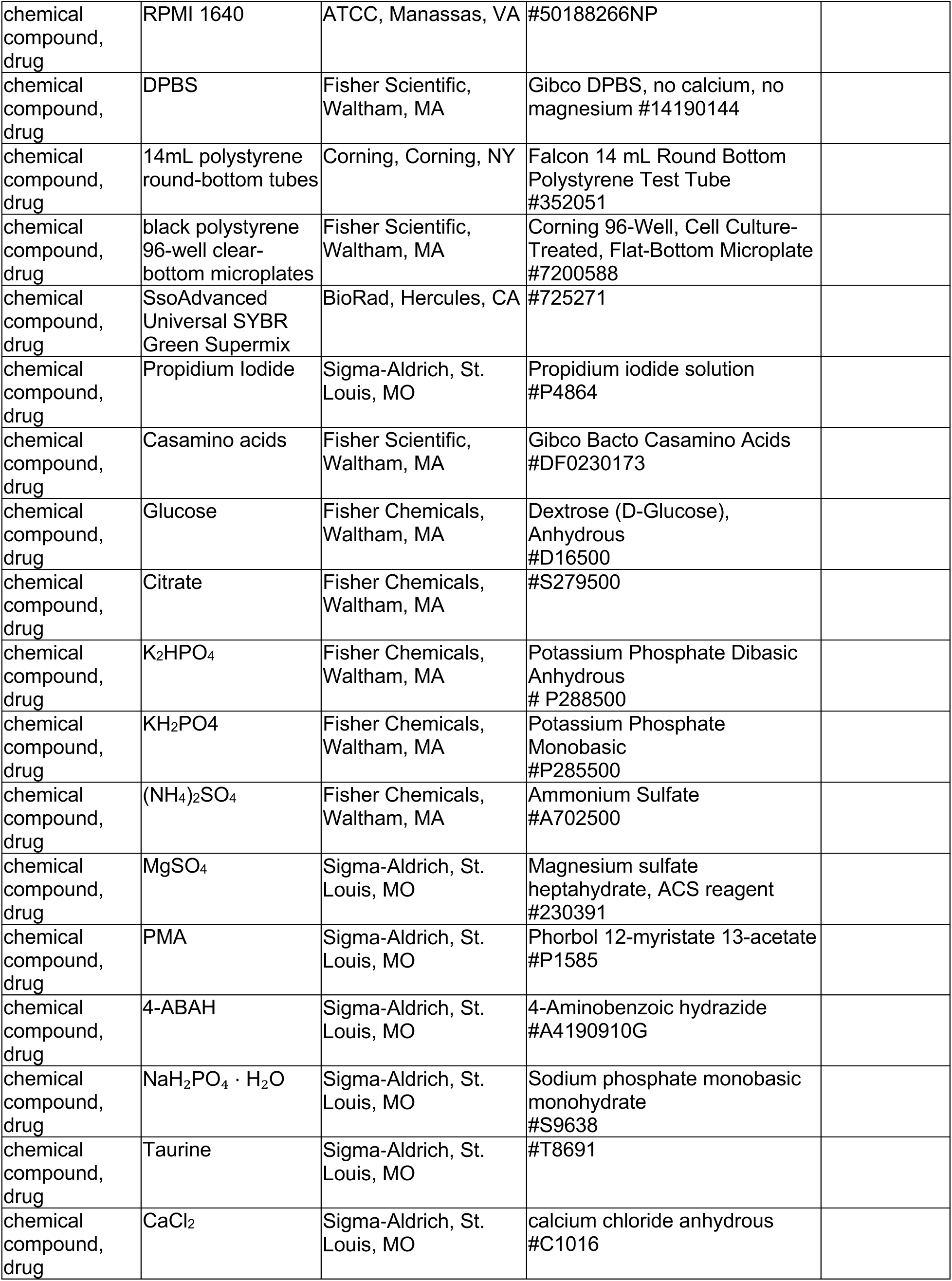
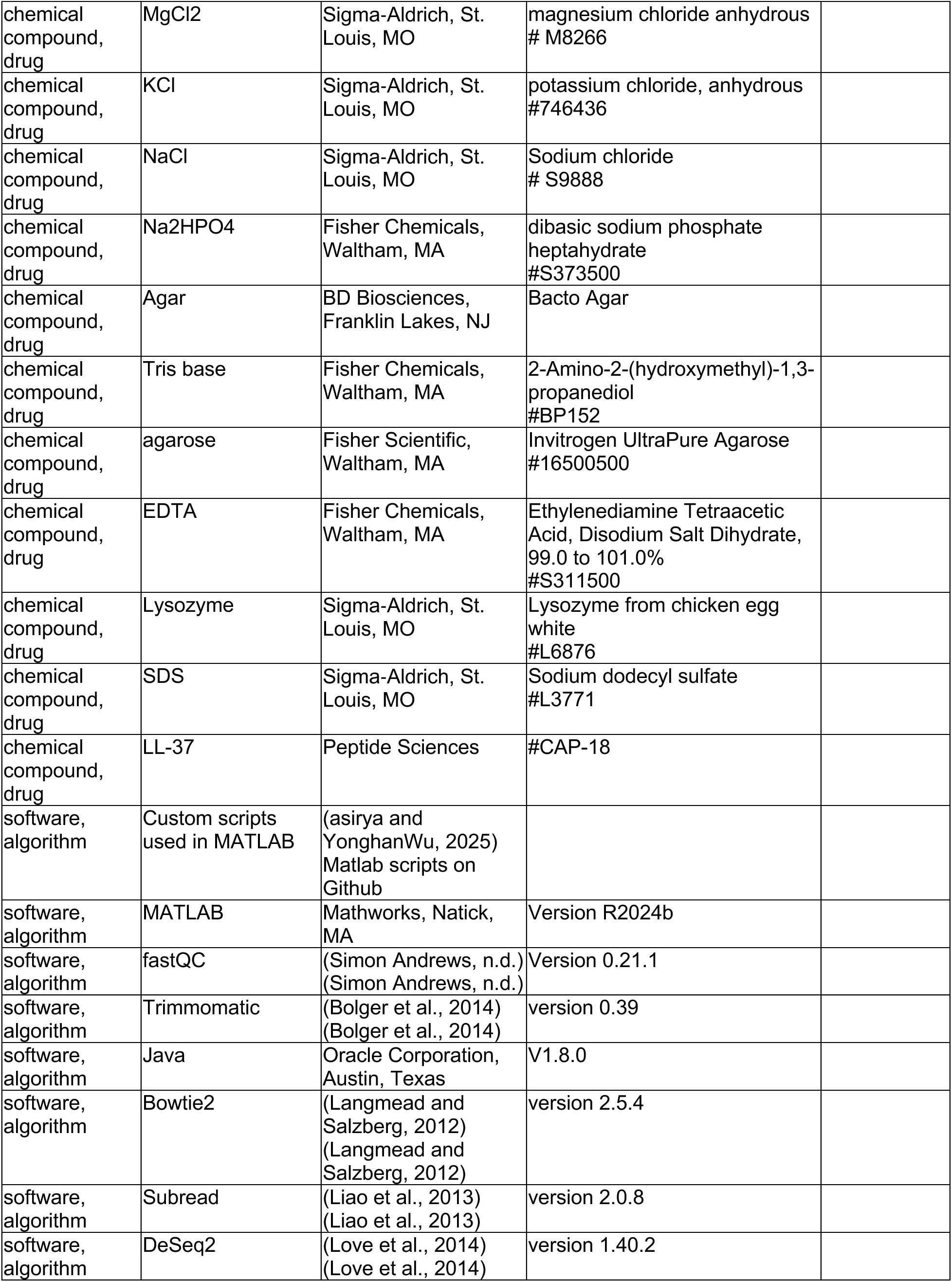
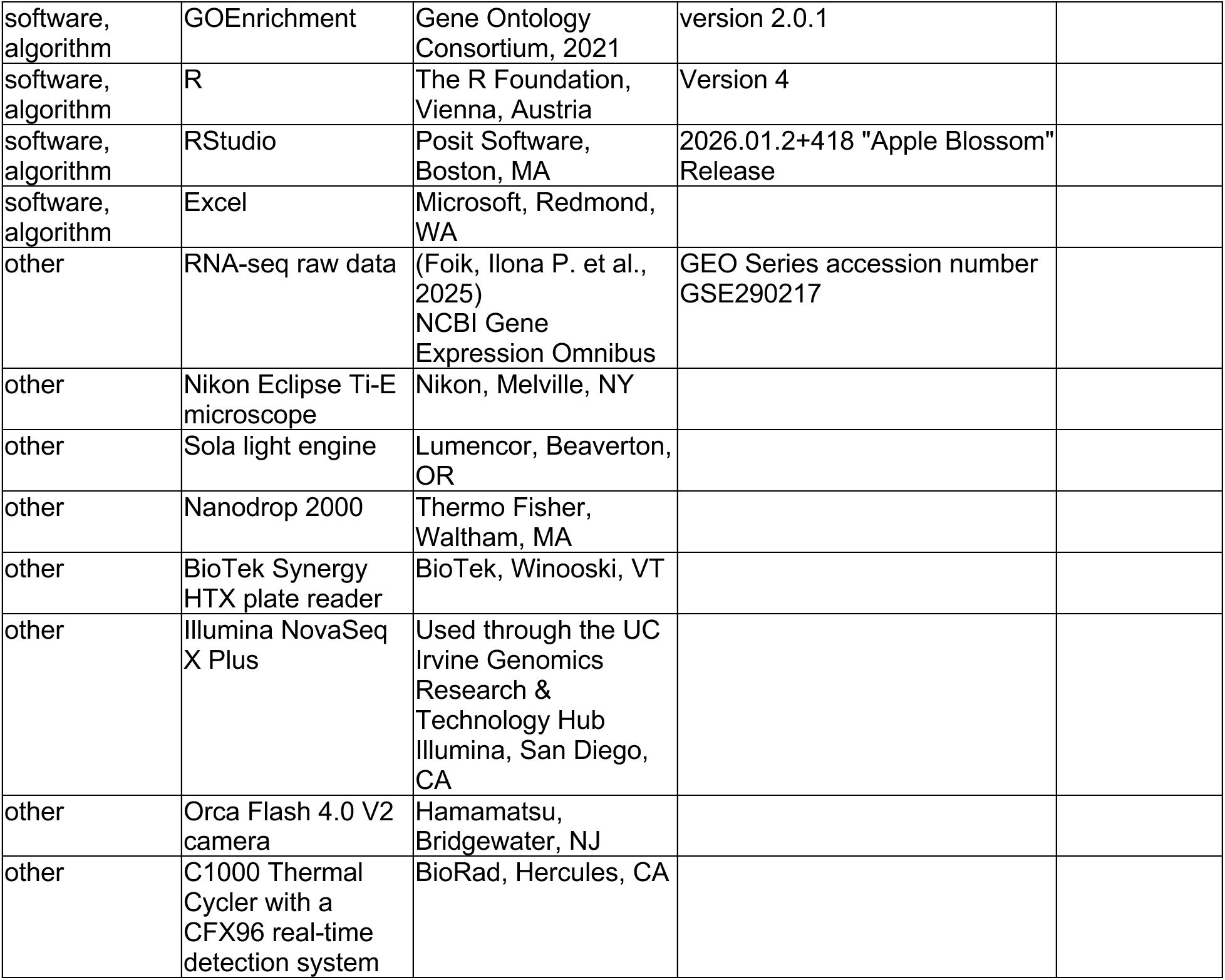

### Bacterial Culture Conditions

Strains were struck out from frozen stocks maintained at -80°C onto plates containing LB-Miller broth and 2% Bacto agar and incubated overnight at 37°C. Single colonies were inoculated into 2mL of modified MinA minimal medium (60.3mM K_2_HPO_4_, 33.0mM KH_2_PO4, 7.6mM (NH_4_)_2_SO_4,_ 1.0mM MgSO_4_; (Malke, 1993) containing 0.2% (w/v) sodium citrate (hereafter referred to as MinA minimal medium), and cultured for 18 hours in a roller drum at 17 rpm and 37 °C.

### Imaging *P. aeruginosa* under flow

Microfluidic devices were fabricated using standard soft lithography techniques as established previously (Siryaporn et al., 2015) with channel dimensions of 100 x 50 μm (width x height). *P. aeruginosa* was cultured for 18 hours in LB-miller medium, diluted 1:100, cultured to mid-exponential phase, and injected into the microfluidic device. Fresh LB was injected at a constant rate of 10μL/min. Images were acquired as described in the fluorescence microscopy section.

### Neutrophil Isolation

Neutrophils were isolated freshly from whole blood as described in (Clark et al., 2018). Briefly, 3% dextran in PBS was used to separate red blood cells (RBCs) from whole blood. Neutrophils were purified from the remaining cells by overlaying on a Ficoll density gradient and centrifugation in for 30 minutes at 500 × g. RBCs were lysed using RBC Lysis Buffer and centrifugation for 5 mins at 300 × g. Supernatant was aspirated and the remaining neutrophil pellets were resuspended in RPMI 1640 medium prior to normalizing cell counts. Neutrophils were used immediately for stimulation, preparation of conditioned media, or co-incubation experiments with *P. aeruginosa* as described below, and purity was assessed in parallel using the fluorescent antibodies APC-CD11b, FITC-CD16 and PE-CD66b and a ACEA NovoCyte Flow Cytometer. This procedure routinely yielded greater than 95% CD11b-positive neutrophil populations.

### Co-incubation of Mammalian Cells with *P. aeruginosa*

Neutrophils were prepared as described above. J774A.1 mouse macrophages were cultured in 10 mL DMEM in T75 flasks at 37 °C with 5% CO_2_. Macrophages were passaged every three days through scraping and were passaged every 5 days through trypsinization. Mammalian cells were washed three times by centrifugation at 300 × g for 5 minutes and resuspension in DPBS, and 1 mL aliquots of 4×10^5^ cells were transferred into flat-bottom dishes. *P. aeruginosa* overnight (∼18h) cultures were centrifuged at 4600 × g for 2 minutes, resuspended in DPBS three times, diluted to a final OD_600_ of 0.2, added to the mammalian cells at a multiplicity of infection of 10, and incubated for 30 minutes at 37°C. Dishes were aspirated until approximately 20 µL of medium remained. 1% agarose pads containing DPBS were placed on top of the samples in the dishes to immobilize bacteria and cells. Samples were imaged as described in the fluorescence microscopy section.

### Conditioned Medium from Neutrophils

To activate neutrophils, 2 million freshly harvested neutrophils per mL in 10mM phosphate buffer (pH 7.4; 2.46 mM monobasic phosphate, 7.54 mM dibasic sodium phosphate) containing 140 mM sodium chloride, 10 mM potassium chloride, 0.5 mM magnesium chloride, 1 mM calcium chloride, 1 mg/mL glucose, and 5 mM taurine (Dypbukt et al., 2005), were treated with 100 ng/mL of PMA in 14mL polystyrene round-bottom Falcon tubes for 1 hour at 37°C and 17 rpm. For MPO inhibition studies, neutrophils were pre-treated with 2 mM of 4-ABAH for 30 mins prior to addition of PMA. This concentration of 4-ABAH was scaled up based on its usage of 500 μM of 4-ABAH to treat one million human neutrophils per mL (Tseng et al., 2018). Supernatant was collected after centrifugation for 5 minutes at 4,600 × g and 24 °C and used as conditioned medium immediately for HOCl detection assay as described below. In parallel, *P. aeruginosa* that were cultured for 18 hours in MinA minimal medium were diluted 1:1000 into conditioned media and were incubated for 3 hours at 37 °C in a rolling drum at 17 rpm and imaged as described in the fluorescence microscopy section.

### HOCl Fluorometric Detection Assay

Conditioned medium was assessed immediately using the Amplite Fluorometric Hypochlorite Assay Kit according to manufacturer protocol. Samples were incubated at room temperature in the dark for 10 minutes and measured for fluorescence using a BioTek Synergy HTX plate with 540/25 nm and 585/10 nm filters for excitation and emission, respectively, and using the autogain set to the 1 μM NaOCl condition. HOCl concentrations were determined by fitting the calibration values to the Hill equation using a three-parameter fit and the lowest value in the data series as the minimum value. HOCl concentrations were background-subtracted by the neutrophil buffer-only sample.

### Treatments of *P. aeruginosa* with oxidizers, antioxidants, and stressors

*P. aeruginosa* were cultured 18 hours in MinA minimal medium and diluted 1:1000 into the same medium. HOCl, H_2_O_2_, HNO_3_, NaOH, NaCl, methionine, L-cysteine, or β-mercaptoethanol (βME) were added to 50 mL cultures at the concentrations indicated in the main text and were incubated in 250 mL flasks for 3 hours at 37 °C in an orbital shaker at 225 rpm. LL-37, histones tobramycin, taurine, and taurine chloramine were added following the 1:1000 dilution and incubated in 14 mL round bottom vented tubes in a roller drum at 17 rpm. For the taurine chloramine experiments, taurine chloramine was prepared by reacting freshly-dissolved NaOCl with taurine in a 1:1 molar ratio as described in (Cunningham et al., 1998). MinA minimal medium lacking ammonium was used as the reaction buffer. All samples were imaged and analyzed as described in the fluorescence microscopy section.

### Fluorescence and Phase Contrast Microscopy

*P. aeruginosa* were immobilized on 1% agarose pads containing MinA minimal medium and imaged immediately. Cultures containing densities below 10 *P. aeruginosa* per frame were concentrated using a syringe filter and imaged immediately. Experiments in Figures 2D and 3F were performed without the syringe filter in order to confirm that the *P. aeruginosa* response was not affected by the filtering procedure.

#### Imaging

Images were acquired using Nikon NIS-Elements software and a Nikon Eclipse Ti-E microscope containing a Nikon 100X Plan Apo (1.45 N.A.) objective, Nikon Ph3 phase contrast condenser annulus, Sola light engine, Hamamatsu Orca Flash 4.0 V2 camera, 459/526/596 dichroic, 575/25 excitation and 641/75 emission filters for mCherry, and 509/22 excitation and 544/24 emission filters for YFP (Semrock, Rochester, NY).

#### Image Analysis

Images were analyzed in MATLAB using custom built software written previously (see the Key Resources Table to download). Briefly, *P. aeruginosa* masks were determined from phase contrast images using an edge-detection algorithm. For size analysis, the total pixel area of each individual *P. aeruginosa* cell was determined by computing the mask area and converting from pixels to μm^2^ by multiplying the mask area by a factor of 0.004225 μm^2^/pixel to account for the microscope camera pixel size and objective magnification. *Fro* expression was quantified by averaging computed ratios of YFP to mCherry fluorescence from the masked areas for at least one hundred individual *P. aeruginosa* cells per condition for each experiment.

### Bacterial Growth Curves

Overnight cultures of bacteria were diluted into MinA minimal medium to an optical density at 600 nm (OD_600_) of 0.01, given treatments, and incubated in sterile 96-well microplates at 37°C in a Synergy HTX multi-mode plate reader with continuous orbital shaking at 180 rpm and a 3mm amplitude, with OD_600_ readings taken every 30 minutes.

### Reverse Transcription Quantitative PCR (RT-qPCR)

For non-murine model samples, bacterial overnight cultures were diluted 1:1000 into fresh MinA medium and incubated for 2.5 hours at 37°C and 225 rpm preceding a 30 minute treatment with 1 µM NaOCl or vehicle control. Samples were concentrated using syringe filters and centrifuged at 13,000 × g for 4 minutes. Pellets were snap-frozen in liquid nitrogen and stored at -80C before performing RNA extraction with the NucleoSpin RNA kit. cDNA was prepared from the using the High-Capacity cDNA Reverse Transcriptase Kit, and quantitative PCR was performed using the SsoAdvanced Universal SYBR Green Supermix using the C1000 Thermal Cycler with a CFX96 real-time detection system. *froA* transcripts were quantified using the primers PA14_froA_v2_F and PA14_froA_v2_R that target PA14_21570 in PA14 and PA3284 in PAO1 (see Key Resources Table). Transcript abundances for each sample were normalized by 5S rRNA abundance using the primers PAO1_PA14_5S_F and PAO1_PA14_5S_R (see Key Resource Table), yielding normalized count thresholds (Cts). The average fold change in transcript abundance was determined by computing the ratio of the averaged normalized Cts and raising by the power of 2, as performed previously (Bru et al., 2019). All primers were validated using a standard calibrating curve to assess Cts and transcript abundance.

### Murine Model of Corneal Infection

Expression of *froA* was assessed in a murine model of corneal infection as described previously (Minns et al., 2023; Sun et al., 2012). The work received prior approval from the Institutional Animal Care and Use Committee at the University of California Irvine under protocol #AUP-21-123. Briefly, *P. aeruginosa* strain PAO1F was cultured to mid-exponential phase in MinA minimal medium that was modified to contain only 0.05% citrate and supplemented with 0.1% casamino acids and 0.2% glucose as the carbon sources, centrifuged, and resuspended in PBS. Corneal epithelia of C57BL/6J mice aged 6-8 weeks were abraded with 3×5 mm scratches using a 25G needle, and 2 µL of PBS containing 5×10^4^ of *P. aeruginosa* strain PAO1F was applied topically. Mice were selected at random and those displaying signs of illness or distress were excluded from the study. Comparable numbers of male and female mice were used in each experimental group. The number of animals used for each experiment was determined based on a power analysis estimate. Mouse eyeballs were collected and homogenized in 1 mL of PBS 0, 2, or 20 hours post-infection and centrifuged for 5 minutes at 150 × g to remove corneal materials. Supernatant was collected and centrifuged for 4 minutes at 13,000 × g. Pellets containing bacteria were suspended in lysis solution (10 mM Tris-HCl, 1 mM EDTA pH 8.0, 0.5 mg/mL lysozyme, 1% SDS) (Bru et al., 2019). RNA was prepared using the NucleoSpin RNA kit and assessed for mRNA expression using RT-qPCR. Samples were not blinded because they were processed as they were collected at different timepoints, which revealed their identity.

### RNA-seq Library Preparation and Data Analysis

*P. aeruginosa* was cultured and RNA was prepared as described in the RT-qPCR section. RNA yield was measured using a Nanodrop 2000. Samples were depleted of ribosomal RNA using the NEBNext rRNA Depletion kit, from which cDNA libraries were constructed using the NEBNext Ultra II Directional Library kit, which were sequenced by the UC Irvine Genomics Research & Technology Hub (Irvine, CA) using an Illumina NovaSeq X Plus using paired-end 150 bp reads at a depth of approximately 10 million reads per sample. Raw reads were checked with fastQC and trimmed and filtered into paired and unpaired reads using Trimmomatic in Java using the ‘PE’ setting. Paired and unpaired reads were aligned to the PA14 genome (NCBI accession NC_008463.1) using Bowtie2 using the default ‘sensitive’ mode, were counted using the featureCounts program in Subread with the ‘countReadPairs’ option enabled for paired-end reads, and were fit into a negative binomial model to compute fold-change in gene expression using DeSeq2. The statistical significance of the changes were computed using the Wald test in DeSeq2. Gene Ontology enrichment analysis for Figure 1A was performed using GOEnrichment.

### Statistical Analyses

Statistical analyses and figures were generated using Excel, R, and RStudio.

### Data, Code, and Materials Availability

Raw data for all figures that contain statistical analyses are available in the supplemental source data files and appendices. All other datasets and software are described in the Key Resources Table.

## ACKNOWLEDGEMENTS

The authors thank Michael Z. Zulu and Stephanie Matsuno for their feedback on prior drafts of this article, Hew Yeng Lai for imaging of bacteria in microfluidic devices, and Hanjuan Shao for assistance with RNA preparation.

## Author Contributions

I.P.F., S.J.K, R.S. and A.S. conceived and designed the experiments. I.P.F., S.J.K., R.S., S.A., M.E.M., L.A.U., L.D., and A.P.H. performed experiments and analyzed data. I.P.F. and L.A.U. wrote the initial draft of the manuscript. S.J.K. and A.S. revised it. All authors edited the paper.

