## Supplementary Materials for "Bacteria detect neutrophils via a system that responds to hypochlorous acid and flow"

Figure Supplements

Supplemental Source Data Titles and Legends

Appendix Tables

#### FIGURE SUPPLEMENTS

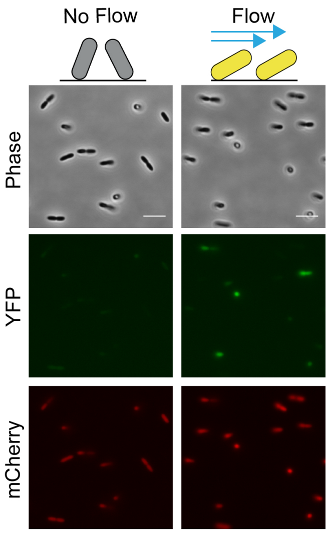


**Figure 1—figure supplement 1. Effect of flow on *fro* expression in *P. aeruginosa*.**
Representative phase contrast and fluorescence images of *P. aeruginosa* under flow (right) or not under flow (left). Fresh LB was flowed at a constant rate of 10 μL/min for 2.5 hours prior to image capture. Scale bars represent 5 μm.

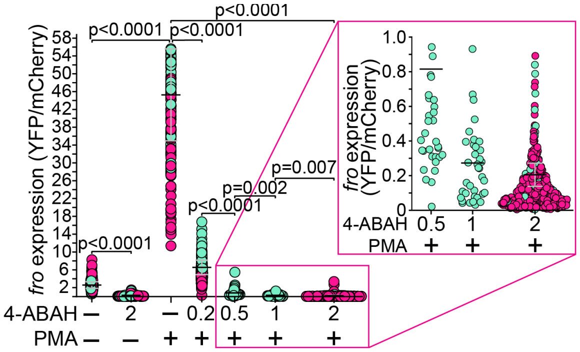


**Figure 2—figure supplement 1. Effects of different 4-ABAH concentrations on *fro* activation.**
*fro* expression as determined by the ratio of YFP to mCherry fluorescence, in *P. aeruginosa* strain AL143 that was cultured for 3 hours in conditioned medium in which neutrophils had been pre-treated with the indicated 4-ABAH concentrations 30 minutes prior to stimulation with 100 ng/mL PMA. Inset shows a zoomed-in view of the conditions indicated. Data points indicate individual *P. aeruginosa*, colors represent independent experiments, bars represent the average of experiments, and grey error bars indicate SEM. P-values were obtained using Welch's t-test on aggregated data from experiments.

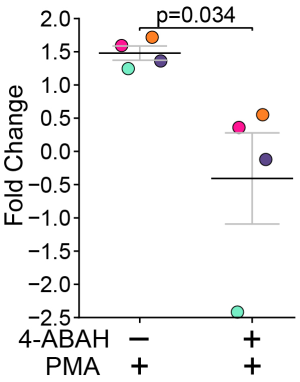


**Figure 2—figure supplement 2. Stimulation of HOCl in neutrophils by PMA and inhibition of activation by 4-ABAH**

Fold-change in HOCl production by neutrophils due to stimulation with 100 ng/mL PMA (activator) compared to unstimulated neutrophils, as measured using a fluorometric hypochlorite detection kit. The fold-repression of HOCl production by PMA-stimulated neutrophils due to pre-treatment with 2 mM 4-ABAH, relative to PMA-stimulation alone was computed using: ($HOCl_{4\text{-ABAH \& PMA-treated}}$ - $HOCl_{unstimulated}$) / ($HOCl_{PMA\ treated}$-$HOCl_{unstimulated}$). An inhibitory value of 1, 0, or a negative value corresponds to no inhibition, total inhibition, or suppression of HOCl below the unstimulated level, respectively. Data points indicate the average from independent experiments utilizing separate neutrophil isolations. Bars indicate the average and errors bars indicate SEM. P-value was obtained using a one-sided Welch's t-test.

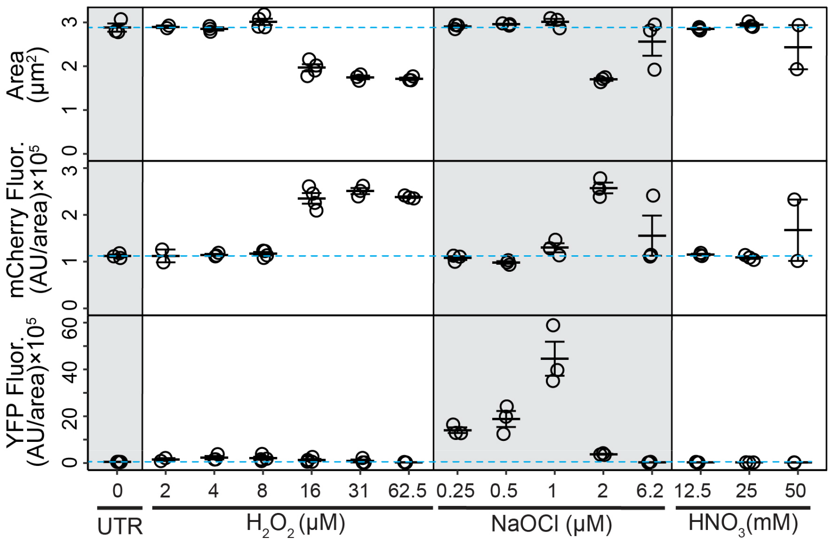


**Figure 3—figure supplement 1. Effects of NaOCl, $H_2O_2$, and $HNO_3$ on cell size and *fro* expression in *P. aeruginosa*.**

*P. aeruginosa* area (top), mCherry fluorescence/area (middle), and YFP fluorescence/area (bottom) of strain AL143 after 3 hours of treatment with $H_2O_2$, NaOCl, $HNO_3$, or vehicle control (UTR). Data points indicate the average from at least one hundred individual *P. aeruginosa* cells, horizontal bars indicate the average of independent experiments, error bars indicate SEM, and blue dashed lines indicate the average of the untreated condition. Gray shading added to improve readability.

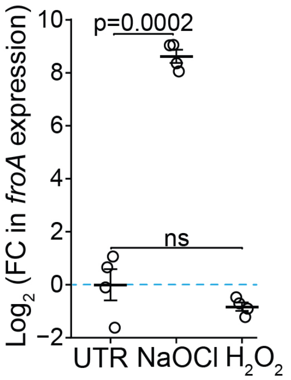


**Figure 4—figure supplement 1. Effect of NaOCl vs $H_2O_2$ on *fro* expression in *P. aeruginosa*.**
Abundance of *froA* transcripts in *P. aeruginosa* strain PAO1F after 30 minutes of treatment with 4 μM $H_2O_2$, 1 μM NaOCl, or no treatment (UTR) as measured by RT-qPCR. Measurements were normalized to 5S ribosomal RNA and the logarithm (base 2) of the fold-change in transcription was computed relative to the untreated condition. $Log_2$(FC) indicates the $log_2$ of the fold-change. Data points represent the average from each experiment, horizontal bars indicate the average of independent experiments (n=4), error bars indicate SEM, and the blue dashed line indicates the average of the untreated condition. P-values were obtained using Welch's t-test, with values of p>0.05 denoted as ns.

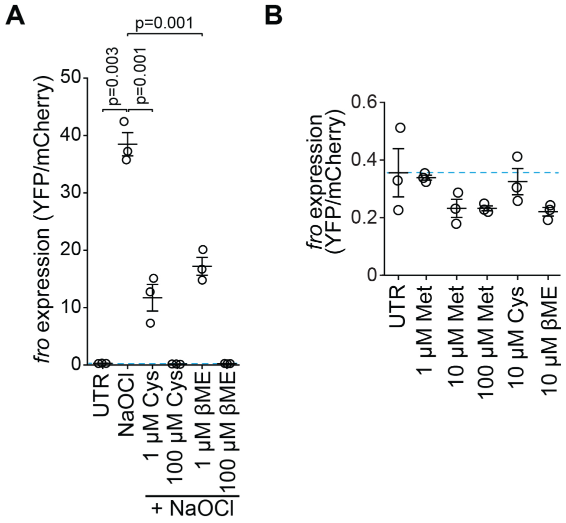


**Figure 5—figure supplement 1. Effects of methionine, cysteine and β-mercaptoethanol on *fro* expression in *P. aeruginosa***
**(A)** *fro* expression as determined by ratios of YFP to mCherry fluorescence in *P. aeruginosa* strain AL143 following no treatment (UTR) or treatment with 1 μM NaOCl alone (replotted from Figure 5B) or combined with the indicated concentrations of cysteine (Cys) or β-mercaptoethanol (βME). **(B)** *fro* expression as determined by ratios of YFP to mCherry fluorescence in *P. aeruginosa* after treatment with Met, Cys, βME, or no treatment. Data points indicate the average from at least one hundred individual P. *aeruginosa* cells, horizontal bars indicate the mean of independent experiments (n=3), error bars indicate SEM, and blue dashed lines indicate the average of the untreated condition. P-values were obtained using Welch's t-test.

#### SUPPLEMENTAL SOURCE DATA FILE TITLES AND LEGENDS

**Figure 1—source data 1**. Source data for heatmap shown in panel A
**Figure 1—source data 2**. Source data for analysis shown in panel D
**Figure 2—source data 1**. Raw image intensity analysis data plotted in panel C
**Figure 2—source data 2**. Raw image intensity analysis data plotted in panel D
**Figure 3—source data 1**. Raw data for growth curves shown in panel A
**Figure 3—source data 2**. Raw image intensity analysis data plotted in panel B
**Figure 3—source data 3**. Raw RT-qPCR data for the plot shown in panel C
**Figure 3—source data 4**. Raw image intensity analysis data plotted in panel D
**Figure 3—source data 5**. Raw image intensity analysis data plotted in panel E
**Figure 3—source data 6**. Raw image intensity analysis data plotted in panel F
**Figure 4—source data 1**. Raw RT-qPCR data for the plot shown
**Figure 5—source data 1**. Raw image intensity analysis data plotted in panel B
**Figure 5—source data 2**. Raw image intensity analysis data plotted in panel C
**Figure 5—source data 3**. Raw image intensity analysis data plotted in panel D
**Figure 6—source data 1**. Raw data for growth curves shown in panel A
**Figure 1—figure supplement 1—source data 1**. Raw image files.
**Figure 2—figure supplement 1—source data 1**. Raw image intensity analysis data plotted
**Figure 2—figure supplement 2—source data 1**. Raw data for computing fold-change in HOCl concentration following three-parameter fits.
**Figure 3—figure supplement 1—source data 1**. Raw image intensity analysis data
**Figure 4—figure supplement 1—source data 1**. Raw RT-qPCR data.
**Figure 5—figure supplement 1—source data 1**. Raw image intensity analysis data plotted in panel A
**Figure 5—figure supplement 1—source data 2**. Raw image intensity analysis data plotted in panel B

### APPENDIX TABLES

| Gene Name | Locus Tag | Description | $Log_2$(Fold-Change) | Fold-Change | p-value |
|---|---|---|---|---|---|
| ***froR***[#] | **PA14_21550** | **RNA polymerase ECF-subfamily sigma-70 factor** | **2.10** | **4.29** | **9.0E-12** |
| ***froA***[#] | **PA14_21570** | **hypothetical protein** | **9.76** | **867.07** | **6.4E-26** |
| ***froB***[#] | **PA14_21580** | **hypothetical protein** | **7.36** | **164.28** | **4.4E-12** |
| ***froC***[#] | **PA14_21590** | **hypothetical protein** | **7.33** | **160.90** | **8.0E-10** |
| ***froD***[#] | **PA14_21600** | **hypothetical protein** | **6.86** | **116.16** | **6.2E-09** |
| ***oprO*** | **PA14_21610** | **pyrophosphate-specific outer membrane porin OprO precursor** | **3.34** | **10.13** | **3.7E-08** |
| ***oprP*** | **PA14_21620** | **phosphate-specific outer membrane porin OprP precursor** | **2.68** | **6.41** | **9.5E-20** |
| ***msrB*** | **PA14_27510** | **methionine sulfoxide reductase B** | **3.59** | **12.04** | **1.6E-14** |
| ***msrQ***[†] | **PA14_62100** | **sulfite oxidase subunit YedZ** | **5.95** | **61.82** | **1.2E-10** |
| ***msrP***[†] | **PA14_62110** | **sulfite oxidase subunit YedY** | **5.18** | **36.25** | **3.0E-11** |
| | **PA14_27520** | **glutathione peroxidase** | **2.71** | **6.54** | **1.4E-08** |
| | **PA14_27530** | **MarR family transcriptional regulator** | **2.02** | **4.06** | **2.7E-07** |
| *bkdB* | PA14_35500 | branched-chain alpha-keto acid dehydrogenase subunit E2 | 1.81 | 3.51 | 7.9E-04 |
| *bkdA2* | PA14_35520 | 2-oxoisovalerate dehydrogenase subunit beta | 1.96 | 3.89 | 7.6E-04 |
| *bkdA1* | PA14_35530 | 2-oxoisovalerate dehydrogenase subunit alpha | 1.88 | 3.68 | 5.8E-04 |
| | PA14_39620 | hypothetical protein | 1.68 | 3.20 | 4.9E-02 |
| *hpd* | PA14_53070 | 4-hydroxyphenylpyruvate dioxygenase | 1.76 | 3.39 | 3.2E-03 |
| | PA14_72360 | hypothetical protein | 1.56 | 2.95 | 2.0E-02 |
| | PA14_72370 | hypothetical protein | 1.61 | 3.05 | 1.1E-02 |
| *mdcC* | PA14_02570 | malonate decarboxylase subunit delta | -1.52 | 0.35 | 2.4E-02 |
| | PA14_32360 | lysR family transcriptional regulator | -1.51 | 0.35 | 2.1E-03 |
| | PA14_32370 | DNA damage-inducible gene | -1.70 | 0.31 | 6.6E-03 |
| *oprN* | PA14_32380 | multidrug efflux outer membrane protein OprN precursor | -3.52 | 0.09 | 4.3E-03 |
| *mexF* | PA14_32390 | RND multidrug efflux transporter MexF | -2.72 | 0.15 | 2.3E-02 |
| | PA14_37270 | LamB/YcsF family protein | -2.60 | 0.16 | 3.6E-02 |
| | PA14_37290 | hypothetical protein | -2.75 | 0.15 | 3.1E-02 |
| | PA14_37310 | hypothetical protein | -2.83 | 0.14 | 1.9E-02 |
| | PA14_37320 | hypothetical protein | -2.39 | 0.19 | 2.7E-02 |
| | PA14_54210 | ATP-dependent protease | -1.89 | 0.27 | 3.8E-02 |
| *groEL* | PA14_57010 | chaperonin GroEL | -2.04 | 0.24 | 1.5E-02 |
| *groES* | PA14_57020 | co-chaperonin GroES | -1.97 | 0.26 | 2.4E-02 |
| *dapB* | PA14_62940 | dihydrodipicolinate reductase | -2.11 | 0.23 | 8.3E-03 |
| *dnaJ* | PA14_62960 | chaperone protein DnaJ | -2.10 | 0.23 | 1.2E-02 |
| *dnaK* | PA14_62970 | molecular chaperone DnaK | -2.34 | 0.20 | 1.6E-02 |
| *grpE* | PA14_62990 | heat shock protein GrpE | -2.00 | 0.25 | 3.1E-02 |

**Appendix 1—table 1. Transcriptional changes due to HOCl stress in WT *P. aeruginosa* vs. Δ*froR* mutant**

Genes that were altered in the wild-type *P. aeruginosa* strain by at least 2.8-fold compared to the Δ*froR* mutant, after 30 minutes of treatment with 1 μM NaOCl and that had p-values less than 0.05. Fold changes were determined from at least three independent experiments and p-values were computed using the Wald test in DeSeq2 (Love et al., 2014, 2014). Annotations are from the Pseudomonas Genome Database (Winsor et al., 2016). Entries are sorted primarily by fold-change and then sub-sorted by gene locus, and **bold** indicates loci that were at least 4-fold upregulated and are displayed in the main text figure. The raw RNA-seq data set is available online (Foik, Ilona P. et al., 2025). [#]The *fro* gene names are labeled based on their usage in (Sanfilippo et al., 2019). [†]The *msrQ* (PA14_62100) and *msrP* (PA14_62110) genes are annotated in the Pseudomonas Genome Database as *yedZ* and *yedY*, respectively, but are referred to here as described in (Juillan-Binard et al., 2017).

| Gene Name | Locus Tag | Description | $Log_2$(Fold-Change) | Fold-Change | p-value |
|---|---|---|---|---|---|
| ***froA***[#] | **PA14_21570** | **hypothetical protein** | **2.92** | **7.57** | **0.0015** |
| ***fpvD***[‡] | **PA14_33550** | **ABC transporter ATP-binding protein** | **2.81** | **7.01** | **0.0000** |
| ***pvdQ*** | **PA14_33820** | **penicillin acylase-related protein** | **2.78** | **6.87** | **0.0008** |
| ***fpvE***[‡] | **PA14_33540** | **ABC transporter permease** | **2.69** | **6.45** | **0.0001** |
| ***pvdA*** | **PA14_33810** | **L-ornithine N5-oxygenase** | **2.67** | **6.36** | **0.0030** |
| | **PA14_33580** | **hypothetical protein** | **2.59** | **6.02** | **0.0001** |
| ***bkdB*** | **PA14_35500** | **branched-chain alpha-keto acid dehydrogenase subunit E2** | **2.49** | **5.62** | **0.0000** |
| ***froC***[#] | **PA14_21590** | **hypothetical protein** | **2.47** | **5.54** | **0.0376** |
| ***froB***[#] | **PA14_21580** | **hypothetical protein** | **2.45** | **5.46** | **0.0211** |
| ***bkdA2*** | **PA14_35520** | **2-oxoisovalerate dehydrogenase subunit beta** | **2.43** | **5.39** | **0.0000** |
| ***fpvC***[‡] | **PA14_33560** | **adhesion protein** | **2.41** | **5.31** | **0.0000** |
| ***bkdA1*** | **PA14_35530** | **2-oxoisovalerate dehydrogenase subunit alpha** | **2.36** | **5.13** | **0.0000** |
| ***froD***[#] | **PA14_21600** | **hypothetical protein** | **2.36** | **5.13** | **0.0444** |
| ***msrQ***[†] | **PA14_62100** | **sulfite oxidase subunit YedZ** | **2.35** | **5.10** | **0.0108** |
| | **PA14_33590** | **hypothetical protein** | **2.28** | **4.86** | **0.0001** |
| ***msrP***[†] | **PA14_62110** | **sulfite oxidase subunit YedY** | **2.25** | **4.76** | **0.0038** |
| ***hpd*** | **PA14_53070** | **4-hydroxyphenylpyruvate dioxygenase** | **2.25** | **4.76** | **0.0002** |
| | **PA14_33570** | **hypothetical protein** | **2.23** | **4.69** | **0.0003** |
| ***msrB*** | **PA14_27510** | **methionine sulfoxide reductase B** | **2.13** | **4.38** | **0.0000** |
| | **PA14_39630** | **hypothetical protein** | **2.07** | **4.20** | **0.0240** |
| | **PA14_52890** | **ring-cleaving dioxygenase** | **2.03** | **4.08** | **0.0099** |
| | PA14_33610 | peptide synthase | 1.99 | 3.97 | 0.0039 |
| *oprO* | PA14_21610 | pyrophosphate-specific outer membrane porin OprO precursor | 1.96 | 3.89 | 0.0011 |
| | PA14_33830 | hypothetical protein | 1.96 | 3.89 | 0.0004 |
| | PA14_33530 | hypothetical protein | 1.96 | 3.89 | 0.0006 |
| *msuD* | PA14_34190 | methanesulfonate sulfonatase MsuD | 1.93 | 3.81 | 0.0000 |
| *oprP* | PA14_21620 | phosphate-specific outer membrane porin OprP precursor | 1.90 | 3.73 | 0.0000 |
| | PA14_72370 | hypothetical protein | 1.87 | 3.66 | 0.0031 |
| *actP* | PA14_22350 | acetate permease | 1.80 | 3.48 | 0.0342 |
| *lipH* | PA14_27090 | lipase chaperone | 1.72 | 3.29 | 0.0006 |
| *pvdG* | PA14_33270 | protein PvdG | 1.71 | 3.27 | 0.0489 |
| *pvdJ* | PA14_33630 | protein PvdJ | 1.71 | 3.27 | 0.0109 |
| | PA14_39620 | hypothetical protein | 1.70 | 3.25 | 0.0472 |
| *msuE* | PA14_34180 | NADH-dependent FMN reductase MsuE | 1.70 | 3.25 | 0.0003 |
| | PA14_72360 | hypothetical protein | 1.65 | 3.14 | 0.0140 |
| *pvdL* | PA14_33280 | peptide synthase | 1.63 | 3.10 | 0.0497 |
| | PA14_34170 | hypothetical protein | 1.58 | 2.99 | 0.0010 |
| | PA14_07870 | ABC transporter substrate-binding protein | 1.57 | 2.97 | 0.0069 |
| | PA14_34260 | ABC transporter permease | 1.56 | 2.95 | 0.0008 |
| | PA14_38640 | CoA transferase subunit B | 1.56 | 2.95 | 0.0044 |
| | PA14_54580 | hypothetical protein | 1.56 | 2.95 | 0.0133 |
| | PA14_33600 | hypothetical protein | 1.56 | 2.95 | 0.0001 |
| | PA14_41790 | hypothetical protein | 1.52 | 2.87 | 0.0020 |
| | PA14_34200 | FMNH2-dependent monooxygenase | 1.52 | 2.87 | 0.0000 |
| | PA14_11790 | amino acid transporter | 1.51 | 2.85 | 0.0090 |
| | PA14_36300 | TetR family transcriptional regulator | -1.50 | 0.35 | 0.0000 |
| | PA14_31580 | acyl-CoA dehydrogenase | -1.55 | 0.34 | 0.0040 |
| | PA14_66350 | acyl-CoA dehydrogenase | -1.61 | 0.33 | 0.0002 |
| *cyoB* | PA14_47190 | cytochrome o ubiquinol oxidase subunit I | -1.76 | 0.30 | 0.0001 |
| | PA14_64310 | ABC transporter ATP-binding protein | -1.80 | 0.29 | 0.0016 |
| | PA14_34520 | ABC transporter permease | -1.80 | 0.29 | 0.0100 |
| *tonB2* | PA14_02490 | hypothetical protein | -1.81 | 0.29 | 0.0001 |
| *cyoA* | PA14_47210 | cytochrome o ubiquinol oxidase subunit II | -1.83 | 0.28 | 0.0000 |
| | PA14_43180 | short chain dehydrogenase | -1.88 | 0.27 | 0.0004 |
| | PA14_09380 | transporter | -1.96 | 0.26 | 0.0263 |
| | PA14_36270 | dehydrogenase | -1.96 | 0.26 | 0.0004 |
| | PA14_36290 | hypothetical protein | -1.98 | 0.25 | 0.0005 |
| | **PA14_53400** | **oxidoreductase** | **-2.01** | **0.25** | **0.0001** |
| | **PA14_36280** | **antibiotic biosynthesis monooxygenase** | **-2.03** | **0.24** | **0.0001** |
| ***cyoE*** | **PA14_47150** | **protoheme IX farnesyltransferase** | **-2.11** | **0.23** | **0.0000** |
| ***cyoC*** | **PA14_47180** | **cytochrome o ubiquinol oxidase subunit III** | **-2.12** | **0.23** | **0.0000** |
| | **PA14_10540** | **iron-sulfur cluster-binding protein** | **-2.12** | **0.23** | **0.0017** |
| ***cyoD*** | **PA14_47160** | **cytochrome o ubiquinol oxidase subunit IV** | **-2.17** | **0.22** | **0.0000** |
| | **PA14_07360** | **hypothetical protein** | **-2.21** | **0.22** | **0.0136** |
| | **PA14_10550** | **sulfite or nitrite reductase** | **-2.30** | **0.20** | **0.0056** |
| | **PA14_34510** | **hypothetical protein** | **-2.47** | **0.18** | **0.0087** |
| | **PA14_34500** | **ABC transporter ATP-binding protein** | **-2.66** | **0.16** | **0.0116** |
| | **PA14_34460** | **hypothetical protein** | **-2.91** | **0.13** | **0.0028** |
| | **PA14_34490** | **hypothetical protein** | **-3.11** | **0.12** | **0.0042** |

**Appendix 1—table 2. Transcriptional changes due to HOCl stress in WT *P. aeruginosa***
Genes that were altered in the wild-type *P. aeruginosa* strain by at least 2.8-fold compared to untreated, after 30 minutes of treatment with 1 μM NaOCl and that had p-values less than 0.05. Fold changes were

determined from at least three independent experiments and p-values were computed using the Wald test in DeSeq2. Annotations are from the Pseudomonas Genome Database (Winsor et al., 2016). Entries are sorted by fold change, and **bold** indicates loci that were displayed in the main text figure. The raw RNA-seq data set is available online (Foik, Ilona P. et al., 2025). [#]The *fro* gene names are labeled based on their usage in (Sanfilippo et al., 2019). [†]The *msrQ* (PA14_62100) and *msrP* (PA14_62110) genes are annotated in the Pseudomonas Genome Database as *yedZ* and *yedY*, respectively, but are referred to here as described in (Juillan-Binard et al., 2017). [‡]The *fpv* gene names are labeled based on their identification as PAO1 orthologs in the Pseudomonas Genome Database

| Gene name | Locus Tag | Description | $Log_2$(Fold-Change) | Fold-Change | p-value |
|---|---|---|---|---|---|
| ***oprN*** | **PA14_32380** | **multidrug efflux outer membrane protein OprN precursor** | **2.90** | **7.46** | **0.0279** |
| ***mexF*** | **PA14_32390** | **RND multidrug efflux transporter MexF** | **2.83** | **7.11** | **0.0271** |
| ***fpvD*** ǂ | **PA14_33550** | **ABC transporter ATP-binding protein** | **2.61** | **6.11** | **0.0002** |
| ***fpvE*** ǂ | **PA14_33540** | **ABC transporter permease** | **2.51** | **5.70** | **0.0004** |
| | **PA14_53830** | **hypothetical protein** | **2.51** | **5.70** | **0.0000** |
| ***fpvC*** ǂ | **PA14_33560** | **adhesion protein** | **2.41** | **5.31** | **0.0001** |
| | **PA14_15435** | **hypothetical protein** | **2.37** | **5.17** | **0.0110** |
| ***mexX**** | **PA14_38395** | **periplasmic multidrug efflux lipoprotein** | **2.25** | **4.76** | **0.0000** |
| ***mexY* (amrB)*** | **PA14_38410** | **efflux protein** | **2.21** | **4.63** | **0.0000** |
| ***merE*** | **PA14_15445** | **mercury resistance protein** | **1.99** | **3.97** | **0.0358** |
| | PA14_34170 | hypothetical protein | 1.86 | 3.63 | 0.0002 |
| | PA14_38140 | glutamine synthetase | 1.81 | 3.51 | 0.0022 |
| | PA14_34200 | FMNH2-dependent monooxygenase | 1.79 | 3.46 | 0.0000 |
| | PA14_38130 | amino acid permease | 1.76 | 3.39 | 0.0001 |
| *msuE* | PA14_34180 | NADH-dependent FMN reductase MsuE | 1.76 | 3.39 | 0.0002 |
| | PA14_33530 | hypothetical protein | 1.75 | 3.36 | 0.0042 |
| | PA14_24360 | hypothetical protein | 1.69 | 3.23 | 0.0000 |
| | PA14_15430 | resolvase, essential for transposition | 1.63 | 3.10 | 0.0025 |
| | PA14_33580 | hypothetical protein | 1.60 | 3.03 | 0.0177 |
| | PA14_32360 | lysR family transcriptional regulator | 1.60 | 3.03 | 0.0022 |
| *cupA5* | PA14_37000 | chaperone CupA5 | 1.59 | 3.01 | 0.0173 |
| | PA14_33570 | hypothetical protein | 1.59 | 3.01 | 0.0135 |
| *merP* | PA14_15470 | periplasmic mercuric ion binding protein, MerP | 1.58 | 2.99 | 0.0259 |
| | PA14_36650 | hypothetical protein | 1.53 | 2.89 | 0.0041 |
| | PA14_36280 | antibiotic biosynthesis monooxygenase | -1.54 | 0.34 | 0.0054 |
| *pscT* | PA14_42640 | translocation protein in type III secretion | -1.54 | 0.34 | 0.0073 |
| *pscO* | PA14_42580 | translocation protein in type III secretion | -1.55 | 0.34 | 0.0184 |
| *cysN* | PA14_57710 | bifunctional sulfate adenylyltransferase subunit 1/adenylylsulfate kinase | -1.55 | 0.34 | 0.0003 |
| *cysW* | PA14_03670 | sulfate transport protein CysW | -1.57 | 0.34 | 0.0013 |
| *cysI* | PA14_40770 | sulfite reductase | -1.58 | 0.33 | 0.0001 |
| | PA14_03710 | hypothetical protein | -1.59 | 0.33 | 0.0002 |
| | PA14_09580 | hypothetical protein | -1.59 | 0.33 | 0.0000 |
| | PA14_42530 | type III secretion protein | -1.63 | 0.32 | 0.0097 |
| | PA14_13000 | transcriptional regulator | -1.65 | 0.32 | 0.0056 |
| *popN* | PA14_42550 | Type III secretion outer membrane protein PopN precursor | -1.67 | 0.31 | 0.0134 |
| *sbp* | PA14_03700 | sulfate-binding protein | -1.67 | 0.31 | 0.0005 |
| *cysT* | PA14_03680 | sulfate transport protein CysT | -1.76 | 0.30 | 0.0015 |
| *pscN* | PA14_42570 | type III secretion system ATPase | -1.77 | 0.29 | 0.0075 |
| | PA14_38920 | hypothetical protein | -1.77 | 0.29 | 0.0000 |
| | PA14_38990 | hypothetical protein | -1.80 | 0.29 | 0.0002 |
| *pcrR* | PA14_42490 | transcriptional regulator protein PcrR | -1.82 | 0.28 | 0.0029 |
| *pscP* | PA14_42600 | translocation protein in type III secretion | -1.84 | 0.28 | 0.0025 |
| | PA14_38970 | two-component sensor | -1.85 | 0.28 | 0.0000 |
| | PA14_27520 | glutathione peroxidase | -1.85 | 0.28 | 0.0003 |
| | PA14_42520 | hypothetical protein | -1.87 | 0.27 | 0.0089 |
| | PA14_36290 | hypothetical protein | -1.87 | 0.27 | 0.0020 |
| | PA14_42630 | translocation protein in type III secretion | -1.89 | 0.27 | 0.0021 |
| | PA14_13010 | hypothetical protein | -1.92 | 0.26 | 0.0008 |
| | PA14_38900 | two-component response regulator | -1.92 | 0.26 | 0.0002 |
| *pscQ* | PA14_42610 | type III secretion system protein | -1.97 | 0.26 | 0.0007 |
| | **PA14_38880** | **hypothetical protein** | **-2.01** | **0.25** | **0.0008** |
| ***exaB*** | **PA14_38850** | **cytochrome c550** | **-2.10** | **0.23** | **0.0008** |
| ***pscR*** | **PA14_42620** | **type III secretion system protein** | **-2.24** | **0.21** | **0.0001** |
| | **PA14_38910** | **sensor kinase** | **-2.30** | **0.20** | **0.0003** |
| ***exaA*** | **PA14_38860** | **quinoprotein alcohol dehydrogenase** | **-2.45** | **0.18** | **0.0002** |

**Appendix 1—table 3. Transcriptional changes due to HOCl stress in the *P. aeruginosa* ΔfroR mutant.**

Genes that were altered in the *P. aeruginosa* Δ*froR* mutant by at least 2.8-fold compared to untreated, after 30 minutes of treatment with 1 μM NaOCl and that had p-values less than 0.05. Fold changes were determined from at least three independent experiments and p-values were computed using the Wald test in DeSeq2. Annotations are from the Pseudomonas Genome Database (Winsor et al., 2016). Entries are sorted approximately by fold change. **Bold** indicates loci that were displayed in the main text figure. The raw RNA-seq data set is available online (Foik, Ilona P. et al., 2025). *The genes *mexX* and *mexY* were identified through BLAST searches against annotated PAO1 genes on Pseudomonas Genome Database. ǂThe *fpv* gene names are labeled based on their identification as PAO1 orthologs in the Pseudomonas Genome Database.

| Anti-σ Locus in PA14 | Anti-σ Locus in PAO1 | Anti-σ Name | Percent of protein containing methionine or cysteine residues | Identified as an anti-σ factor in PA14 by: | Reference(s) |
|---|---|---|---|---|---|
| PA14_20000 | PA3409 | *hasS* | 0.60 | Ortholog of *hasS* in PAO1 | (Dent et al., 2019) |
| PA14_01860 | PA0150 | | 0.90 | Ortholog of PA0150 in PAO1 | (Llamas et al., 2008) |
| PA14_06170 | PA0471 | *fiuR* | 0.92 | Ortholog of *fuiR* in PAO1 | (Llamas et al., 2006) |
| PA14_37980 | PA2051 | | 0.94 | Ortholog of PA2051 in PAO1 | (Llamas et al., 2008) |
| PA14_46650 | PA1364 | | 1.06 | Otero-Asman et al., 2019 | (Otero-Asman et al., 2019b) |
| PA14_13450 | PA3900 | *fecR* | 1.26 | Ortholog to *fecR* in PAO1 | (Llamas et al., 2008) |
| PA14_47390 | PA1301 | *hxuR* | 1.52 | Ortholog of *hxuR* in PAO1 | (Otero-Asman et al., 2019a, 2019b) |
| PA14_26610 | PA2895 | *sbrR* | 1.65 | McGuffie et al., 2016 & Pseudomonas Genome Database | (McGuffie et al., 2016) |
| PA14_33780 | PA2388 | *fpvR* | 1.80 | Ortholog of *fpvR* in PAO1 | (Rédly and Poole, 2003) |
| PA14_32720 | PA2467 | *foxR* | 1.82 | Ortholog of *foxR* in PAO1 | (Llamas et al., 2006) |
| PA14_20730 | PA3351 | *flgM* | 1.85 | Ortholog of *flgM* in PAO1 & Pseudomonas Genome Database | (Dasgupta et al., 2003) |
| PA14_28980◊ | N/A | | 2.19 | Chevalier et al., 2019 | (Chevalier et al., 2019; Otero-Asman et al., 2019b) |
| PA14_64690 | PA4895 | | 2.33 | Ortholog of PA4895 in PAO1 | (Llamas et al., 2008) |
| PA14_37420 | PA2094 | | 2.49 | Ortholog of PA2094 in PAO1 | (Llamas et al., 2008) |
| PA14_55540 | PA0676 | *vreR* | 2.80 | Ortholog of *vreR* in PAO1 | (Llamas et al., 2009) |
| PA14_54420 | PA0763 | *mucA* | 3.08 | Pseudomonas Genome Database | (Wu et al., 2004) |
| PA14_39810 | PA1911 | *femR* | 3.13 | Ortholog of *femR* in PAO1 | (Llamas et al., 2008) |
| PA14_46820 | PA1350 | | 3.59 | Ortholog of PA1350 in PAO1 | (Potvin et al., 2008) |
| PA14_69390 | PA5255 | *algQ* | 4.35 | Pseudomonas Genome Database | (Ambrosi et al., 2005) |
| PA14_41610 | PA1774 | *cfrX* | 5.95 | Ortholog of *cfrX* in PAO1 | (Chevalier et al., 2019; Otero-Asman et al., 2019b) |
| PA14_21560◊ | N/A | *froI*# | 7.59 | Boechat et al., 2013 | (Boechat et al., 2013) |

**Appendix 2—table 1. Abundance of methionine and cysteine residues in *P. aeruginosa* anti-σ factors.**

The percentage of methionine and cysteines for anti-σ loci were determined using the *P. aeruginosa* strain PA14 sequence. Loci in PA14 were identified previously or determined through ortholog identification using the Pseudomonas Genome Database (Winsor et al., 2016) based on the PAO1 gene name or locus that were reported previously. Entries are sorted by the ‘Percentage of protein containing methionine or cysteine residues’ column. #The *froI* gene name is based on its usage in (Sanfilippo et al., 2019). ◊These PA14 anti-σ factors are not found in PAO1.
